# Towards interoperable modeling of toehold-mediated strand exchange circuits across DNA nanotechnology and engineering biology

**DOI:** 10.64898/2026.07.22.739222

**Authors:** Zoila Jurado, Geoffrey Taghon, Samuel W. Schaffter

## Abstract

Originally developed for DNA nanotechnology, toehold-mediated strand exchange (TMSE) circuits are gaining traction in synthetic biology due to their high programmability, seamless integration with biological components, and robust operation across diverse environments and cell types. However, while forward-engineering in synthetic biology has benefited from automated genetic circuit modeling pipelines, there is currently a lack of accessible, automated tools for the mechanistic modeling of TMSE circuits integrated with these systems, hindering the development of new biotechnologies. The TMSE-BioCRNpyler Library allows TMSE molecules to be transcribed RNAs or fixed-concentration nucleic acids while leveraging existing BioCRNpyler features, such as upstream transcription regulation and downstream gene regulation. We demonstrate this library’s applicability by modeling published applications of TMSE circuits spanning a wide range of applications, including simple *in vitro* reactions, cell-free biosensors, and *in vivo* microbial and mammalian systems. Additionally, we validated that models compiled using the TMSE-BioCRNpyler Library produced results with < 0.2 % relative error compared to multiple models of TMSE previously developed in the literature. Finally, to streamline interoperability with existing models, we developed txt2biocrnpyler. This accompanying tool converts chemical reaction networks from the literature into a BioCRNpyler-ready source script and a Systems Biology Markup Language XML file — a standard data format for sharing and simulating biological models. The TMSE-BioCRNpyler Library serves as a powerful new resource for the rational, automated design of molecular information processing systems.

## Introduction

A major goal of engineering biology is the ability to use first principles to rationally design genetic circuits that process molecular information in biological systems. Paramount to this goal are mathematical models that simulate the dynamics of the biomolecular species that make up these engineered systems [1, 2, 3]. Great strides have been made in this direction, with genetic circuit modeling and design software packages gaining broader adoption and accuracy, particularly for transcription-factor-based genetic circuits [4, 5]. These models are broadening our understanding of these systems by implicating undesigned secondary reactions [6], in addition to guiding and accelerating how we design genetic circuits to achieve a desired outcome. Thus, models play an integral role in the Design-Build-Test-Learn (DBTL) cycle of engineering biology [7].

Beyond genetic circuits, *i.e*., circuits that rely on gene expression to process and transduce information, enzyme-free circuits composed of fixed concentrations of nucleic acids that process information via toehold-mediated strand exchange (TMSE) reactions [8, 9] offer an attractive medium for programming biology. TMSE circuits rely entirely on nucleic acid base pairing interactions, making them easy to rationally program [10, 11, 12], and versatile for use across different application spaces [13]. Initially demonstrated in non-biological contexts, TMSE-based logic is increasingly being incorporated with genetic circuit components. For example, TMSE circuits have been combined with transcription-based biosensors [14], transfected into mammalian cells [15, 16] and genetically encoded [17, 18, 19] to regulate gene expression in both cell-free protein expression systems and bacteria [20, 21]. Despite the increasing utilization of TMSE in engineering biology, we currently lack a satisfactory modeling package that supports flexible integration of TMSE reactions with genetic circuitry. Existing modeling packages are either tailored towards the non-biological context of the DNA nanotechnology and computing [22, 23] or the transcription factor-based genetic circuits of the engineering biology community [4, 5]. Previous attempts at creating modeling packages that bridge these two fields either failed to gain broad adoption or are no longer actively managed [22, 24, 25].

BioCRNpyler [26] is an actively managed Python package for generating chemical reaction networks (CRNs) that has been primarily leveraged for modeling genetic circuits in cell-free and cellular settings [27, 28, 29, 30]. BioCRNpyler contains a predefined library of biochemical classes and modeling parameters for common engineering biology systems, while also providing sufficient flexibility to introduce additional reactions and tune parameters. Here we developed a TMSE-specific library in BioCRNpyler, TMSE-BioCRNpyler Library, a suite of software tools to support the compilation and simulation of TMSE reactions alongside genetic circuits in BioCRNpyler (Figure 1). The TMSE modules we developed allow TMSE components and mechanisms to be easily implemented in, and incorporated with, the existing capabilities of BioCRNpyler. To demonstrate the utility and flexibility of our TMSE modules, we compiled and simulated more than ten published examples, successfully integrating TMSE circuit models with upstream transcriptional processes and downstream gene expression pathways. These models can be applied to systems containing both DNA and RNA molecules and systems operating in both cell-free and cellular settings.

**Figure 1:**
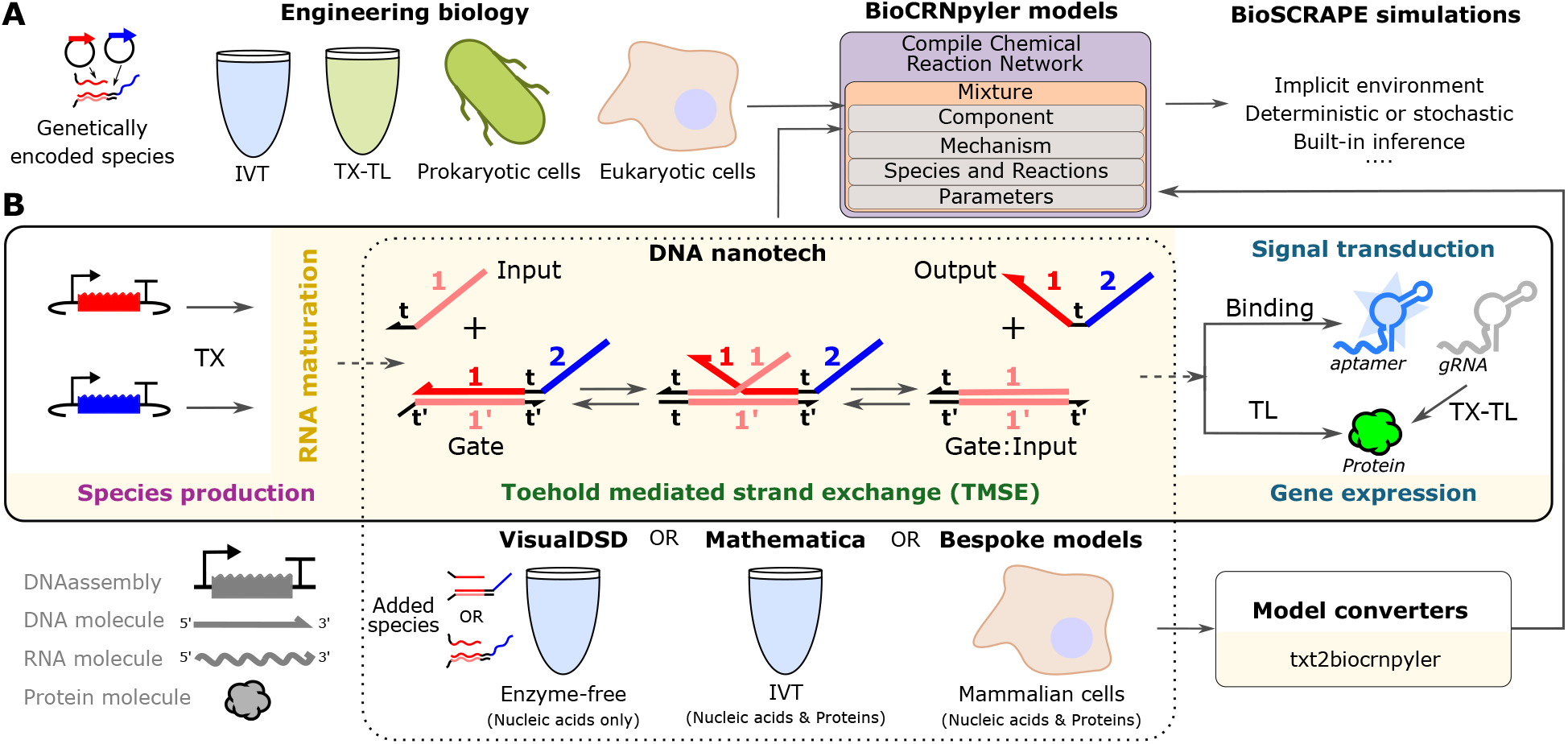
Schematic of the BioCRNpyler modeling workflow, TMSE circuits, and current modeling platforms that span engineering biology and DNA nanotechnology. **(A)** Overview of engineering biology application models compiled in BioCRNpyler and simulated in Bioscrape. **(B)** Overview of the TMSE-BioCRNpyler Library. Sections highlighted in yellow represent new components and mechanisms added to BioCRNpyler to model TMSE circuits (see Supplementary Information Section 2). We introduce txt2biocrnpyler, a standalone tool that expands upon standard SBML converters [33] by translating text-based CRNs directly into BioCRNpyler-ready Python scripts. The dotted box in the middle of the panel illustrates TMSE reactions in DNA nanotechnology applications, in which DNA molecules are added directly to the reaction environment, rather than being expressed. Although the molecules are depicted as DNA, the same TMSE reactions apply to RNA, whether it is added directly or expressed *in situ*. In the reaction network schematics throughout this work, DNA and RNA molecules are designated with straight and wavy lines, respectively.

The compiled BioCRNpyler models can be simulated using Bioscrape — a complementary Python package for flexible CRN simulation [31] — or exported in Systems Biology Markup Language (SBML), the XML-based standard for sharing and storing computational models of biological processes [32]. Furthermore, Bioscrape can import and extend SBML models generated by BioCRNpyler and other standard platforms, including VisualDSD [22] for TMSE circuits or iBioSim [5] for genetic circuits.

However, many published models are not provided in SBML format, but rather described as lists of reactions or differential equations in supporting information, making it onerous to import and modify them in BioCRNpyler. Further, expanding models directly in Bioscrape is much more limited than expanding them within BioCRNpyler. To enable the reuse and repurposing of existing models within BioCRNpyler and Bioscrape, we developed txt2biocrnpyler — a Python-based translator that converts text-based lists of chemical reaction equations into a BioCRNpyler-ready script or an SBML version 3 XML file. Using txt2biocrnpyler, we demonstrate the conversion, extension, and simulation of model descriptions from the literature within BioCRNpyler and Bioscrape. Together these results illustrate how the TMSE-BioCRNpyler Library and txt2biocrnpyler tool can bridge modeling across DNA nanotechnology, DNA computing, and engineering biology to rapidly advance new cross-field applications (Figure S1).

## Methodology

### Development and features of TMSE components and mechanisms in BioCRNpyler

TMSE reactions typically involve two types of nucleic acid molecules (Figure 1 and Figure S2), single-stranded Inputs and partially double-stranded Gate complexes composed of two strands. Gate molecules contain both an input domain (designated as a red *1* in Figure 1) and an output domain (designated as a blue *2* in Figure 1). Each of these domains contains a short region known as a toehold (designated as a black *t* in Figure 1). The toehold of the input domain on a Gate can recruit a sequence-complementary Input (designated by a matching input domain *1* ) to bind to the Gate and initiate a process of branch migration in which the Input and Output strands stochastically migrate back and forth on the Gate until the Output strand is released. Upon release the Output has a newly exposed toehold that allows it to initiate downstream strand exchange reactions with Gates that possess a complementary input domain (domain *2* in Figure 1). Thus, the input and output domains of the strand exchange molecules dictate which molecules can react with one another. Strand exchange is often designed to be reversible such that an Output strand can reinitiate branch migration with complementary Gate:Input complexes to drive the Input back off. However, the reaction is typically driven towards Output production by using an excess of Input or coupling the final Output of the circuit to an irreversible reaction (Reporter Gates in Figure S2) Beyond Inputs, Gates, and Outputs, TMSE circuits often use other nucleic acid molecules that serve different functions but have the same general features of Input and Gate molecules. These include for example, Fuel strands, Threshold Gates, THErr Gates, and Reporter Gates (Figure S2).

To enable straightforward modeling of TMSE circuits in BioCRNpyler, we began by developing TMSE-specific component and mechanism modules (Tables S1-S3). Within BioCRNpyler, components create species with defined attributes and mechanisms generate mass action reactions between species. Components and mechanisms are fed into mixtures, which provide BioCRNpyler context regarding the environment in which the reactions take place. In principle, TMSE reactions can be added individually to BioCRNpyler using generic components and mechanisms, such as `Reaction.from_massaction`. However, TMSE circuits can comprise more than 100 distinct molecules and contain myriad different connections between molecules [10, 11, 12], making the number of class instantiations unwieldy (Table S4). Therefore, it is preferable for the user to define initial TMSE molecules and allow TMSE-specific components to identify how the strands will interact and generate reactions and products for the final model. Thus, we designed the input to the TMSE-specific components as a Python dictionary in which each key maps to a list specifying the strand exchange molecule and its connectivity (domains) to other strand exchange molecules. In addition to reducing the number of class instantiations, structuring the code this way mimics the inputs to experiments and reduces the chances of introducing errors when compiling a new model.

Strand exchange molecules are defined using input dictionaries with the following structure:

~~~
{‘molecule1’: [sd_type, domain_i, domain_j],
 ‘molecule2’: [sd_type, domain_i],
  …}.
~~~

For example, keys can match experimental nomenclature, which may be more detailed than strand exchange molecule definitions in the model. The dictionary values are lists that define the strand exchange molecule, `sd_type`. The string `sd_type` specifies the functional identity of the strand exchange molecule (i.e. ‘Input’, ‘Output’, ‘Gate’, ‘TGate’ for Threshold Gates, ‘Reporter’, ‘THErr’ for Toehold exchange riboregulator, or ‘Fuel’). The next two entries of the list, ‘domain_i’ and ‘domain_j’, can be integers or strings that define the identities of the input and output domains of the strand exchange molecule (Figure S2), thereby dictating the interactions between molecules. In the case that only one domain is provided, e.g. for an Input, the strand exchange molecules generated will assume identical domains for *i* and *j*. In the case of THErr molecule, ‘domain_j’ will be a list composed of two elements denoted below,

~~~
{‘therr_mol’: [‘THErr’, domain_i, [ domain_j1, domain_CDS ]]},
~~~

where the ‘domain_CDS’ denotes a protein coding sequence.

We developed the `StrandExchangeMol()` component to automatically generate DNA and RNA molecules capable of strand exchange, with input and output domains explicitly defined as attributes, directly from the strand exchange input dictionary. If the system is composed of only added DNA or RNA molecules at fixed concentrations, `StrandExchangeMol()` is initialized with the `material_type` parameter set to ‘dna’ or ‘rna’, respectively. However, if the system incorporates RNA molecules dynamically generated via transcription, the transcription parameter must be enabled in `StrandExchangeMol()`. When this option is enabled, `transcription = True`, DNA templates for transcription and their corresponding RNA molecules are automatically created in the model. Crucially, a separate component must be defined for each distinct set of processes the molecules undergo. For example, if a system contains static DNA, transcribed RNA, and static RNA strand exchange molecules, these distinct groups must be instantiated in separate `StrandExchangeMol()` calls.

To simulate systems with transcribed RNAs, a transcription mechanism must be introduced (as depicted in the “Produce” block in Figure 2A). The `StrandExchangeMol()` component is fully compatible with existing transcription mechanisms in BioCRNpyler, including `SimpleTranscription()` and `Transcription_MM()`, and can also be coupled to custom, user-defined transcription models. Additionally, `StrandExchangeMol()` seamlessly integrates into various pre-defined mixtures, as illustrated later in Figure 4. This modularity allows users to easily tailor the biological context and level of abstraction to match their experimental conditions.

**Figure 2:**
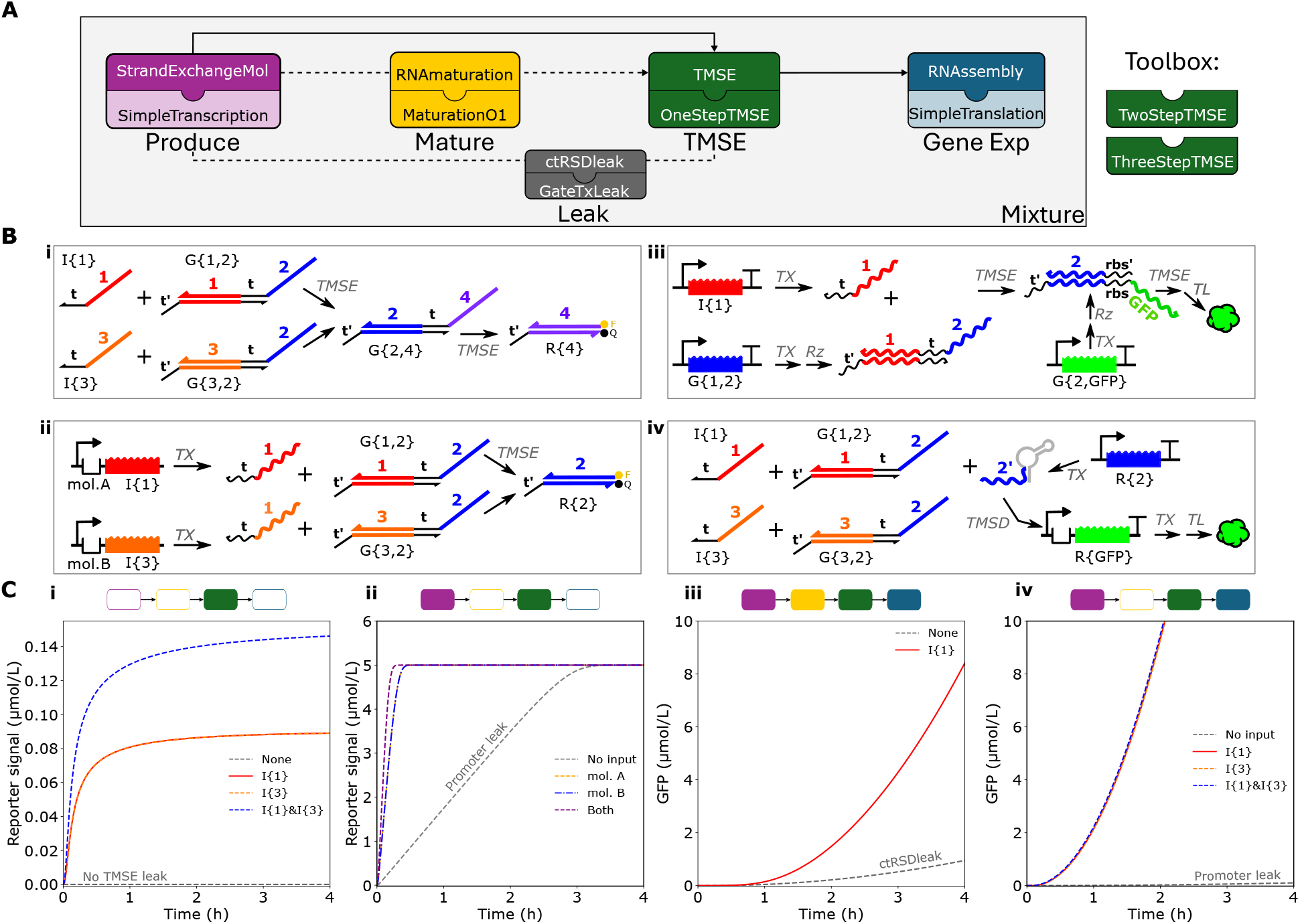
Comprehensive overview of the design, components, and simulation of TMSE circuits in BioCRN-pyler. **(A)** Illustration of primary TMSE modules, including their respective interactions and connections, with dashed lines indicating optional pathways. Each component (upper block) is highlighted, along with the corresponding mechanism (lower block). Additional mechanisms are displayed in the toolbox. **(B)** Schematics of the four distinct TMSE circuit types we modeled in this study: **(i)** DNA seesaw circuits, **(ii)** transcription-factor-based biosensors, **(iii)** genetically encoded RNA-based TMSE circuit, and **(iv)** DNA-based TMSE circuit designed to regulate gene expression in mammalian cells. In the schematics, DNA and RNA TMSE components are designated with straight lines and wavy lines, respectively. Explicitly named species represent the chemical components directly added to the reaction mixture, corresponding to the model inputs passed to `StrandExchangeMol()`. **(C)** The respective simulated results for each of the four circuits shown in (B). Colored blocks above the plots indicate which TMSE-specific modules from (A) were used to create the models for each simulation.

The heart of the TMSE-specific modules are the `TMSE()` component and `OneStepTMSE()` mechanism (green block in Figure 2A). `TMSE()` takes in a concatenated list of BioCRNpyler species, created through `StrandExchangeMol()` and creates a dictionary of all the relevant TMSE reactants and products between these strand exchange molecules based on their domains, effectively abstracting the reaction generation process. The reaction dictionary returned by `TMSE()`, is then used to compile reversible mass action reactions using one of three TMSE mechanisms. The `OneStepTMSE()` mechanism models TMSE as a single reversible bimolecular reaction between reactants and products, ignoring the process of branch migration. The `TwoStepTMSE()` and `ThreeStepTMSE()` [8] mechanism introduces an intermediate step, shown in Figure 1, providing a layer of additional detail required to capture situations when dissociation of the Output RNA is rate-limiting. By default, all TMSE reactions are modeled as reversible. However, if branch migration or Output RNA dissociation is rate-limiting, resulting in a k_rev_ rate constant of zero, the mechanism automatically implements an irreversible, forward-only reaction. For modeling enzyme-free, DNA-only TMSE circuits — where all strand exchange molecules are DNA added at fixed concentrations — the model input dictionaries can be fed directly to `TMSE()` to build the reaction dictionary. For systems containing a mixture of DNA and RNA TMSE molecules it is recommended to create all species through `StrandExchangeMol()` and feed those to `TMSE()` to identify interactions and create the remaining reaction species.

Species from `StrandExchangeMol()` can be fed directly into `TMSE()`. Alternatively, if RNA maturation is required we introduced an optional component for RNA maturation between `StrandExchangeMol()` and `TMSE()`. Maturation of RNAs into functional products through cleavage, splicing, capping, and folding are common in biology [34], and are often utilized in synthetic settings [35, 36]. RNA maturation is particularly important for cotranscriptionally encoded RNA strand displacement (ctRSD) circuits, which use a self-cleaving ribozyme to create double-stranded RNA gates suitable for TMSE [17, 18, 19]. To include a maturation step on an RNA molecule, the maturation parameter must be set to True within `StrandExchangeMol()`, `rna_maturation = True`. This will designate the molecules as preprocessed (“pre”) RNAs that cannot interact with other molecules. These pre-RNA molecules are fed to the `RNAmaturation()` component, which uses, a first-order reaction mechanism `MaturationO1()` to generate processed RNA molecules that can now interact with other molecules. These processed RNA molecules can then be fed to `TMSE()` alongside other strand exchange molecules. The output of the TMSE circuit can be used as-is for direct signal plotting, manually coupled to auxiliary reactions to model signal transduction pathways (as depicted in Figure 1B), or connected to a translation mechanism to drive gene expression. To connect the outputs of TMSE circuits to protein expression, we created a new component to complete the Gene Exp block (Figure 2A). BioCRNpyler already contains mechanisms for modeling RNA translation, but to connect these mechanisms to products from TMSE reactions we needed a component that specifies which RNAs can be translated. This connection is often done with an existing BioCRNpyler component called `DNAassembly()`, which defines a DNA template that encodes for the transcription and translation of a protein. However, `DNAassembly()` is not set up to handle the input and output domains required for specifying TMSE reactions and would result in duplicate DNA species. Thus, to define an RNA assembly for translation consisting of a ribosome binding site, transcript, and protein, we wrote the `RNAassembly()` component — analogous to the `DNAassembly()` component. RNAs generated from the previous components can be assigned as the transcript for translation in the `RNAassembly()` component. When compiled with existing translation mechanisms in BioCRNpyler, the specific named protein will be translated from the specified transcript.

The components we described thus far assume ideal behavior and do not account for the undesired ‘leak’ reactions known to occur in many TMSE circuits. To illustrate how to account for unintended transcription of Output RNAs, we developed a dedicated leak component. Specifically, we modeled a type of transcriptional leak frequently observed in cotranscriptionally encoded RNA strand displacement (ctRSD) circuits [18]. To include this type of leak, the leak variable must be set to ‘ctRSDleak’ within `StrandExchangeMol()`, `leak =` ‘`ctRSDleak`’. When leak is enabled, `StrandExchangeMol()` will generate two RNA species — Gate and Gate_dummy_ — transcribed from the same DNA for all Gate molecules. This component facilitates models of leaky gene expression by employing the `GateTxLeak()` mechanism, which generates reactions that produce an Output RNA with domains identical to the parent Gate DNA. The leak reaction is modeled according to the following stoichiometric scheme:

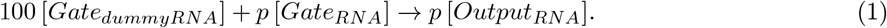

Here, the Gate_dummyRNA_ acts as a non-reactive counter representing the total RNA transcribed, while Gate_RNA_ represents the direct RNA produced by parent Gate DNA. In this formulation, the unintended production of Output_RNA_ is governed by the baseline transcription rate constant (k_tx_) and is proportional to defined leak percentage (*p*). This approach treats the leak as a cotranscriptional or RNA folding error. Consequently, the DNA does not transcribe two distinct RNA molecules, but rather a specific percentage (*p*) of the total RNA produced inherently results in the leaky Output RNA. Crucially, this formulation enforces mass conservation. Mechanisms that model other forms of TMSE leak reactions [37] could be added in a similar manner.

To compile a final TMSE CRN, we use a mixture in BioCRNpyler to unite all the components, mechanisms, parameters, species, and reactions, as highlighted in Figure 1. A mixture in BioCRNpyler provides context of the model’s environment in addition to tying components to mechanisms and their respective parameters (reaction rate constants, concentrations, etc). The BioCRNpyler mixture library contains mixtures for Cell-like or Extract-like environments that model transcription and translation at different levels of mechanistic detail to more accurately model the experimental environment. Finally, the CRN is compiled from the defined mixture and either simulated using Bioscrape or exported in SBML format for publication or use in other simulators compared in [38].

### Importing published models into BioCRNpyler using txt2biocrnpyler

Many models described in the engineering biology literature, rather than being provided in SBML format, are described as lists of reactions or differential equations, requiring new BioCRNpyler and Bioscrape users to rebuild these models from scratch. To circumvent this issue, we developed a Python-based translator tool, txt2biocrnpyler. As its name suggests, the tool accepts a text-based list of chemical reactions (see Supplementary Information Section 4.2) and outputs BioCRNpyler-ready scripts. These scripts can then be modified and extended using existing BioCRNpyler packages, or simulated directly in Bioscrape. The tool, txt2biocrnpyler, can also directly export SBML version 3 XML files that can be shared or simulated with any SBML simulator, such as Bioscrape. This tool adds a new capability (generation of BioCRNpyler-ready Python scripts) to the existing suite of tools for converting between different modeling paradigms and SBML [33]. Briefly, a deterministic algorithm extracts species, reactions, kinetic rate laws, stoichiometries, compartments, and parameter values from natural language or lists of mathematical equations.

To ensure model integrity, txt2biocrnpyler validates both output pathways through distinct mechanisms. When generating BioCRNpyler-ready scripts, an automatic validation check occurs because the model is compiled prior to generating the final SBML output. Conversely, the direct-to-SBML path validates outputs against an SBML version 3 Pydantic schema before serialization via libsbml. The full txt2biocrnpyler source code is available at https://github.com/usnistgov/txt2biocrnpyler, and can be installed on any system with a Python interpreter. We provide both an txt2biocrnpyler GUI web application and a Jupyter notebook for intuitive, non-command-line use. A step-by-step tutorial for installing, launching, and using the browser-based txt2biocrnpyler graphical user interface to generate these BioCRNpyler-ready Python scripts, alongside a flowchart detailing the tool workflow, is provided in the Supplementary Information Section 4.

With the advent of large language models (LLMs), it is natural to ask whether a deterministic, algorithm-based tool like txt2biocrnpyler is necessary, or if LLMs can directly translate literature into BioCRNpyler scripts. We argue that dedicated algorithms remain essential for two primary reasons: data security and strict reliability and reproducibility. First, tools like txt2biocrnpyler process data entirely on local hardware, protecting sensitive intellectual property and preventing the exposure of proprietary designs to third-party cloud servers. Second, and more importantly, automated workflows require absolute syntactic consistency — standard LLMs struggle to meet. To benchmark the LLM reliability and reproducibility, we tasked three prominent LLMs (Gemini, ChatGPT, and Claude) with converting the chemical reaction equations of AND gate from the literature [14] into a functional BioCRNpyler script. Running the same zero-shot prompt 10 times per model revealed significant variance in performance (Table S5).

While Claude achieved a perfect 10/10 success rate (occasionally providing the raw SBML XML alongside the Python script), Gemini produced functional scripts in only 6 out of 10 attempts, and ChatGPT succeeded in just 4 out of 10. Noticeably, the LLMs suffered from both outright failure modes — such as syntax hallucinations that crashed the code — and non-deterministic output variations, like inconsistent species naming. Even when scripts were functionally executable, unpredictable naming conventions required manual verification to ensure species mapped correctly. By replacing non-deterministic text generation with a deterministic algorithm, txt2biocrnpyler eliminates these bottlenecks, ensuring secure, reproducible, and fully automatable code generation.

## Results and Discussion

### Benchmarking and Validation of TMSE-BioCRNpyler

To validate the TMSE-specific components and mechanisms we developed worked correctly, we compared the simulation results of our compiled models with those of other TMSE models in the literature. To quantitatively compare simulation results, we computed the reciprocal relative error between the two results across relevant species in the simulation [38] (Supplementary Information Section 5). The goal of this analysis is to reveal if there are any issues in how our TMSE-specific modules handle model assembly. As some of the simulation results we compared used different numerical solvers, we expected some discrepancies between results. However, if these discrepancies are small with a < 1 % relative error, then the differences are likely due to the ODE solvers rather than a difference in the compiled models themselves.

Conducting this analysis for two different test cases: a mixture of RNA- and DNA-based circuits [14] (Figure S4 to S5) and cotranscriptionally encoded RNA strand displacement (ctRSD) circuits [19] (Figures S6 to S8) — revealed < 0.15 % relative error between results from models produced with the TMSE-BioCRNpyler vs existing literature software (Table 1, Supplementary Information Section 5).

**Table 1:** Benchmarking TMSE-BioCRNpyler’s accuracy against literature models. A comparison of the maximum relative error between models compiled using the TMSE-BioCRNpyler library and their respective original published results.

| Model | Circuit | Relative error |
| --- | --- | --- |
| ctRSD-sim | 2-layer | < 0.08 % |
| ctRSD-sim | Fan-Out | < 0.10 % |
| ctRSD-sim | Fan-In-Out | < 0.12 % |
| Jung2022 | OR | < $1.0 \times 10^{-4}$ % |

### Simulating literature examples of integrated TMSE and genetic circuits

Having validated the underlying modules of the TMSE-BioCRNpyler Library, we sought to illustrate the broader utility by using the library to model TMSE. TMSE circuits have been used in a broad range of biologically relevant environments and applications [13], and this breadth of use motivated our development of TMSE modules for BioCRNpyler. Nucleic-acid-only TMSE circuits are often operated in a pH-buffered solution containing cations (enzyme-free conditions). In these conditions, the kinetic parameters for TMSE reactions as a function of toehold length and sequence have been extensively characterized and used for predictive modeling [8, 9]. Both DNA- and RNA-based TMSE circuits have been coupled to *in vitro* transcription (IVT) reactions, which has enabled the development of complex dynamic systems that can perform sophisticated information processing tasks. Further, TMSE circuits have been used to regulate protein expression in cell-free transcription-translation conditions (cell-free TX-TL) and extract-based transcription-translation conditions (extract TX-TL) [20]. In addition to test tube environments, genetically encoded TMSE circuits have been operated in bacteria [21] and transfected into mammalian cells (cellular conditions) to control cellular behavior [15, 16]. The systems we model in Figure 2 use the TMSE-BioCRNpyler Library to instantiate all the examples described above, while leveraging existing BioCRNpyler capabilities, such as simulating different environments and leveraging pre-defined gene regulation, TX, and TL models.

In enzyme-free conditions, DNA-based circuits that operate entirely on TMSE have been scaled up to hundreds of molecules to execute digital calculations, molecular pattern recognition, and supervised learning [10, 11, 12]. The development of these massive circuits was made possible by predictive modeling. Several computational tools — such as VisualDSD and the Seesaw Compiler — have been developed to design and simulate DNA-based TMSE circuits [22, 23]. Although these models can theoretically be extended to incorporate genetic circuit interactions, substantial barriers still hinder their broader adoption, particularly among experimentalists. Expansions to a model either require intimate knowledge of the software or require integrating multiple computational tools [22, 24, 25], some of which are no longer supported. Thus, we first illustrate how to use the TMSE modules we developed to implement enzyme-free DNA-based circuits in BioCRNpyler.

To simulate a DNA-based circuit that implements OR logic (illustrated in Figure 2B(i)) after defining the DNA molecules in a dictionary, `TMSE()` can be called directly. `TMSE()` checks which strand exchange molecules can react based on their domains and returns a dictionary of strand exchange reactants and products. The component and the `OneStepTMSE()` mechanism are then used as mixture parameters compiled into a CRN and simulated in Bioscrape (Figure 2C(i)). Different levels of model detail can be achieved using `TwoStepTMSE()` or `ThreeStepTMSE()` as demonstrated in Figure S9.

A fundamental part of DNA circuits is a ‘seesaw’ element, which uses a set of reversible TMSE reactions to amplify and propagate signals across circuit layers. When used in conjunction with a Threshold Gate, different types of logical operations can be achieved, as shown in the simulations presented in Figure S10. These examples show how to build DNA-based TMSE CRNs from scratch, but in many cases the models for large circuits have already been developed and it would be beneficial to be able to import those into Bioscrape and extend as needed. Both VisualDSD and the Seesaw Compiler tools can export models in SBML format, which can then be imported into Bioscrape and directly simulated. Figure S11 demonstrates simulation of a DNA-based circuit, composed of 520 molecules, that executes cellular automata transition functions; the SBML formatted file for this simulation was taken from [23].

A straightforward route for connecting TMSE information processing circuits to non-nucleic acid inputs is through introduction of transcriptional regulation. For example, allosteric transcription factors that sense small molecules or ions to regulate transcription can convert the presence of a molecule of interest into expression of RNA inputs that can then be processed with TMSE reactions [14]. However, adding the regulated transcription reactions to existing software tools specifically developed for DNA-based TMSE [22, 23] is non-trivial. Likewise, common software for simulating transcription-factor-based genetic circuits [4, 5] do not accommodate TMSE reactions. As a result, to incorporate transcription and DNA-based TMSE circuits, models are typically developed from scratch [14].

Figure 2B(ii) illustrates an example from the literature that combines transcription and DNA-based TMSE reactions to execute OR logic on molecules A and B that regulate transcription. To simulate this circuit in BioCRNpyler, the two Input RNA molecules are defined in `StrandExchangeMol()` with the transcription parameter set to True then used in `TMSE()`. To automatically build the reactions for transcription-factor-based regulation of input expression — including ‘leaky’ expression as described in [14], we use the existing `RepressiblePromoter()` component in BioCRNpyler. This allows us to build and simulate a variety of the circuits developed in [14] in just a few lines of code (Figure 2C(ii) and Figure S12). The model we present here differs slightly from the model described in [14], which explicitly models transcription factors and RNA polymerase binding and unbinding.

However, our txt2biocrnpyler tool can convert the chemical reactions described in the Supporting Information of [14] into a BioCRNpyler script that can be simulated or extended to incorporate new reactions (Figure 3). This workflow, illustrated in Figure 3A, allows users to seamlessly build upon the originally published model (Figure 3B), iteratively appending new biological mechanisms, such as translation (Figure 3C) and LacI-mediated transcriptional control, ultimately evolving the base model into a functional NIMPLY circuit (Figure 3D) [14, 39]. The same model additions can be done within Bioscrape using an SBML model as the input (see `SI_NOTgate_Bioscrape.ipynb` in the associated GitHub repository), however, working in BioCRNpyler makes certain expansions easier, such as adding translation by simply calling the `SimpleTranslation()` mechanisms or adding degradation by simply changing the mixture.

**Figure 3:**
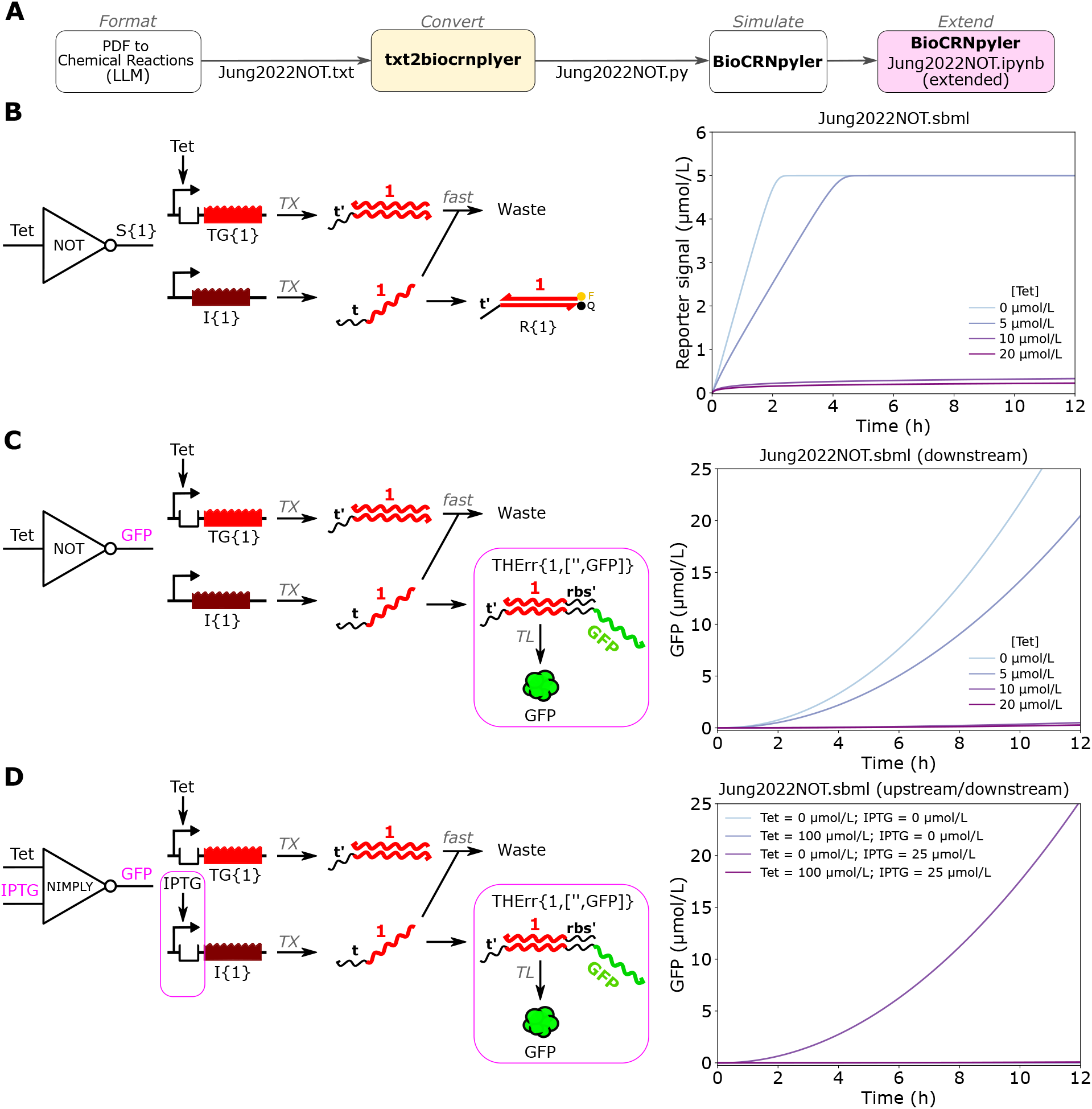
Using txt2biocrnpyler and Bioscrape to simulate and extend models from the literature. **(A)** A model for the NOT circuit from the Supplementary Information of [14] was converted into SBML using the txt2biocrnpyler tool and then simulated or extended to include additional reactions in Bioscrape. **(B)** Bioscrape simulation results of the original model after conversion to SBML [14]. **(C)** Extension of (B) that adds downstream reactions to the model. The DNA reporter was replaced with a toehold exchange riboregulator and a translation mechanism was added. **(D)** Extension of (C) that adds upstream reactions to the model. The I{1} DNA template was modified to conduct LacI-mediated transcriptional regulation resulting in a NIMPLY circuit similar to one previously developed in ROSALIND [14, 39]. Explicitly named species represent the chemical components directly added to the reaction mixture, corresponding to the model inputs passed to `StrandExchangeMol()`.

**Figure 4:**
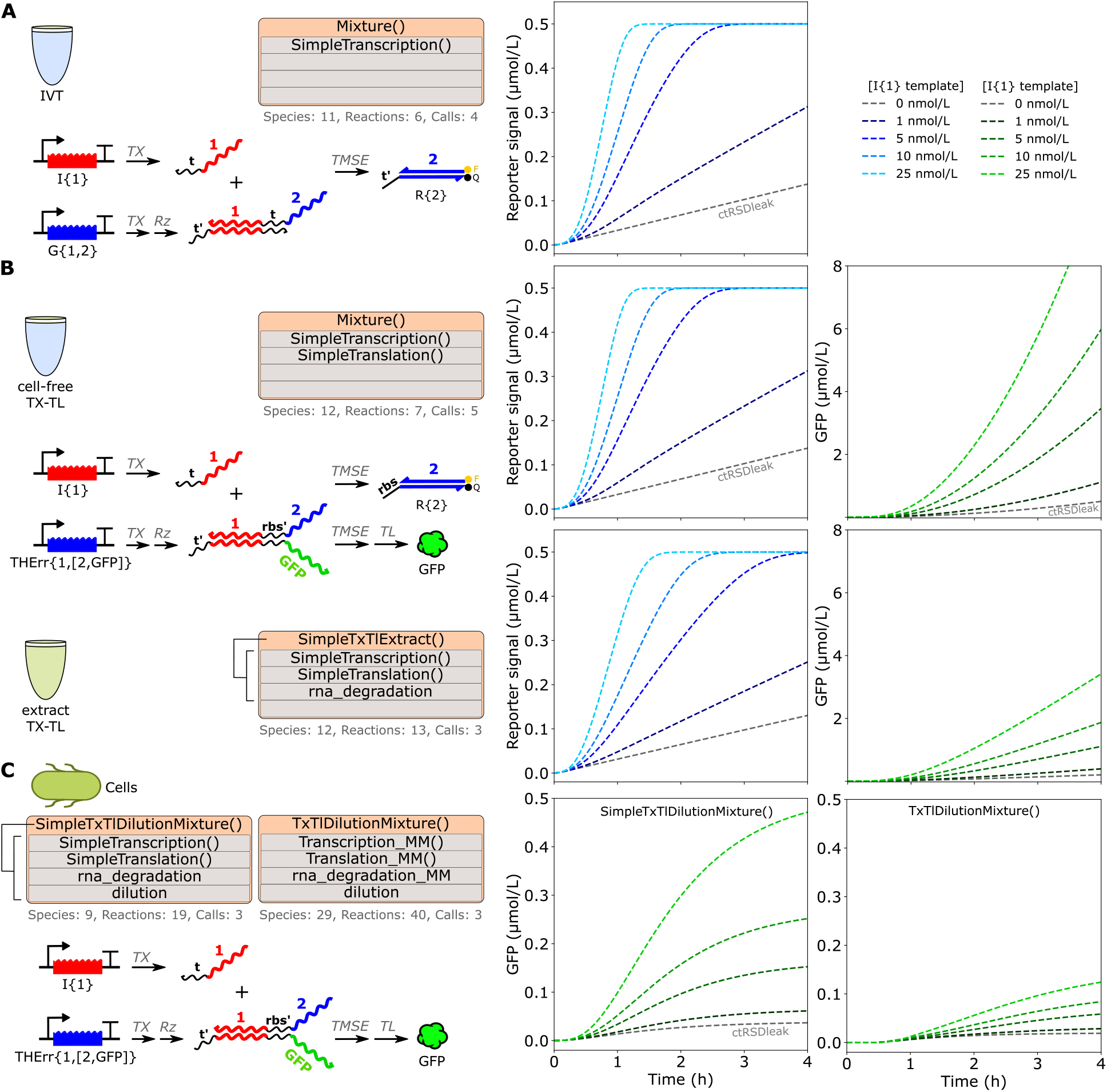
Using the TMSE-BioCRNpyler Library to simulate ctRSD circuits across environments. **(A**,**B)** Schematics and simulations for a ctRSD circuit in (A) IVT conditions [18] and **(B)** cell-free TX-TL conditions (top) or extract TX-TL conditions (bottom) [20]. For the cell-free TX-TL simulation, the `SimpleTranscription()` and `SimpleTranslation()` mechanisms were called directly for use in the final mixture. For the TX-TL simulations, a pre-defined mixture `SimpleTxTlExtract()` was used, which calls the `SimpleTranscription()` and `SimpleTranslation()` mechanisms and adds an RNA degradation mechanism. **(C)** Schematic and simulation results for a ctRSD circuit in cells [21]. Here, the pre-defined mixtures `SimpleTxTlDilutionMixture()` or `TxTlDilutionMixture()` were used, which call transcription and translation mechanisms, add an RNA degradation mechanism, and add a global dilution term for cell division. `TxTlDilutionMixture()` uses Michaelis-Menten kinetics for enzymatic reactions. All simulations also used the `MaturationO1()`, `OneStepTMSE()`, `GateTXLeak()` mechanisms developed in this study. The number of species, reactions and mechanism calls for each model are shown below the mixture and mechanism blocks. Explicitly named molecules are directly added to the reaction mixture.

A promising new direction for TMSE circuits is the development of versions composed entirely of RNA molecules that can be genetically encoded for continuous operation in cells. These cotranscriptionally encoded RNA strand displacement (ctRSD) circuits use self-cleaving ribozymes to generate dsRNA complexes suitable for strand exchange [17, 18, 19]. Further, ctRSD gates have recently been adapted into toehold exchange riboregulators that connect TMSE to the regulation of protein expression [20, 21]. While a software package simulating arbitrary ctRSD circuits in IVT environments exists [40], expanding this software to include new reactions — such as translation — is not straightforward. Further, the matrix implementation used to build and simulate the CRNs is prohibitively slow for large circuits.

Figure 2B(iii) illustrates a ctRSD circuit that regulates protein expression in cell-free TX-TL environments. In this case to simulate this ctRSD circuit, all DNA molecules are transcribed and the two Gates are passed through the `MaturationO1()` mechanism to include the first-order reaction associated with ribozyme cleavage. `RNAassembly()` is used to specify which Gate(s) can be translated to produce a protein and the resulting assembly is used by a translation mechanism. To compile and simulate the model (Figure 2C(iii)), we chose a simple translation mechanism (RNA →Protein) that mimics a nuclease-free PURExpress environment by assuming no RNA or protein degradation. While this specific example includes translation, ctRSD circuits can also be simulated under strictly IVT conditions and integrated with DNA-based TMSE components (Figure 4A).

As all molecules of ctRSD circuits are genetically encoded, they can easily be tested in cell-free or cellular conditions, as has recently been reported in *E. coli* -based extract and *E. coli* [20, 21]. Simulating the circuits in these new application spaces using BioCRNpyler is seamless, simply requiring the selection of an appro-priate `Mixture()` from the available options. Using an alternative mixture such as `SimpleTxTlExtract()` maintains simple TX-TL reactions but automatically adds degradation to all RNA molecules (Figure 4B, bottom). Changing mixture to `SimpleTxTlDilutionMixture()` includes TX-TL reactions, RNA degradation reactions, and includes a dilution reaction to account for cell division (Figure 4C, left). Model complexity can also be easily extended to use Michaelis-Menten kinetics for all enzymatic reactions — thereby doubling the number of species and reactions considered due to the creation of intermediate species and reactions — by changing the `Mixture()` in a single line of code (Figure 4C, right).

In addition to using genetically encoded TMSE circuits in bacteria, fixed concentrations of TMSE components have been transfected into mammalian cells and used to process information and regulate gene expression [15, 16]. The modeling of such systems can take multiple abstractions from a well-mixed solution to a compartmentalized environment — more closely resembling the experimental setup — which can be easily implemented using BioCRNpyler and our TMSE modules. Figure 2B(iv) shows a system in which TMSE is used to regulate the activity of a CRISPR guide RNA, which in turn activates gene expression. To model this system, we begin by using `StrandExchangeMol()` to define the initial RNA and DNA molecules. Then we call on `TMSE()` to check for strand exchange interactions between molecules. To implement CRISPR-Cas systems from the literature [16], we define two additional species Cas9 and CRISPR, and a reaction for the activation of the CRISPR-Cas system by the Signal RNA from the TMSE reactions. The activated CRISPR-Cas system is then used to define an `ActivatablePromoter()`, which serves as the promoter for a `DNAassembly()` component that expresses GFP.

The results in Figure 2C(iv) do not account for spatial compartmentalization, such as the physical separation of transcription in the nucleus and translation in the cytoplasm typical of eukaryotic cells [16]. To introduce compartmentalization more analogous to the actual experiment [16], we can use additional BioCRNpyler modules for membrane proteins [41]. In Figure S13, we show how the TMSE modules can be used conjunctively with the membrane components to incorporate spatial aspects into the model, accounting for the spatial separation of transcription and translation in mammalian cells.

In addition to supporting transcription and translation enzymatic reactions, BioCRNpyler has mechanisms for any type of catalysis that can be added. For example, Figure S14 illustrates how to include a specific RNase in a model of a mixed nucleic acid system [42]. Catalysis can be modeled with differing levels of complexity, from a simple first-order reaction to Michaelis-Menten enzyme kinetics.

## Conclusion

To bridge modeling needs across DNA nanotechnology and engineering biology, we developed the TMSE-BioCRNpyler Library, comprising BioCRNpyler modules designed specifically for compiling TMSE reactions. The TMSE modules allow for the customization of components, mechanisms, and parameters of desired TMSE reactions. Implementation in BioCRNpyler enables the integration of TMSE reactions with genetic circuitry, facilitating the design, analysis, and predictive modeling of complex biochemical networks. To demonstrate the adaptability of the TMSE-BioCRNpyler Library, we assessed the modules through the reconstruction and simulation of over ten published TMSE applications. Additionally, to enable interoperability between existing published models and BioCRNpyler, we developed txt2biocrnpyler to convert models described as chemical reaction lists into BioCRNpyler-ready scripts or SBML Level 3 XML files. By providing a common language and framework for modeling TMSE circuits across environments, we envision accelerating the development of innovative engineering biology applications of nucleic acid computing.

A key advantage of our approach is that we can leverage existing features of BioCRNpyler and Bioscrape to extend the complexity of the models with simple function calls. For example, we illustrate how to incorporate our TMSE-specific modules with existing BioCRNpyler components and mechanisms to model transcriptional regulation (Figure 2), model circuits across different environments (Figure 4), and model compartmentalization (Figure S13). Although most of our examples focused on simple models of TX and TL, our suite of TMSE-specific modules are compatible with more detailed models of these processes that have already been developed or are currently in development [41, 43, 44]. Incorporating other mechanisms for transcriptional regulation to encompass other biotechnologies that have interfaced with TMSE circuits [45, 46, 47] should also be straightforward in BioCRNpyler. BioCRNpyler has a growing library of pre-defined parameters for many common genetic circuit reactions, and our modules could be extended to populate rate constants based on properties of TMSE components (i.e. length, sequence, binding energy, etc.) [8, 9]. Along these lines, while we only used Bioscrape to simulate the compiled TMSE examples, the parameter inference features could be leveraged to parameterize TMSE reactions in cell-free and cellular environments [48, 49].

Furthermore, with the txt2biocrnpyler tool, we have simplified the transition to modeling with BioCRNpyler and Bioscrape without having to manually reconstruct existing models from scratch. In many cases this conversion is a “drag and drop” process, which should increase adoption of the SBML format and enable sharing and integration of models currently sequestered in publications and supplemental materials (Supplementary Information Section 4). One remaining challenge for integrating models from different publications or fields is reconciling different naming conventions for the species and rate parameters.

While our current TMSE modules offer broad flexibility, future iterations would benefit from the inclusion of several additional features. For example, additional modules that cover other common leak reactions [37] or other undesired reactions like toehold occlusion [50] would be beneficial. The current TMSE modules are designed to keep track of molecules with two domains, but expanding to cover molecules with multiple inputs and outputs would broaden capabilities [18, 51]. Additionally, other types of TMSE reactions [52] could be added, such as cooperative hybridization [53, 11, 54] or the use of handholds [55].

Our work provides a template for how other molecular programming paradigms could be incorporated into BioCRNpyler to interface with models of genetic circuits. In fact, many other systems rely on bimolecular reactions between molecules with specific input and output domains. For example, strand exchange and strand displacement have also been used to regulate DNA and RNA polymerization reactions [56, 57, 58, 59, 60], which could be encompassed in future modules. Further, other types of bimolecular transcription regulation mechanisms, such as those described in Refs. [46, 61, 62], could be encompassed as interfacing components in future modules. Modules that handle multi-domain tracking could also be utilized to encompass protein-based CRNs inspired by TMSE circuits [63, 64]. Further, many riboregulators rely on strand displacement opening up an RNA hairpin to regulate gene expression [65, 45, 47]. Our current modules are flexible enough to be modified to simulate such systems. For example, hairpin-based riboregulators could be modeled using the `TwoStepTMSE()` or `ThreeStepTMSE()` mechanisms with an Output dissociation rate constant set to 0. The resulting TMSE intermediate can then be fed to `RNAassembly()` for translation.

More broadly this work highlights the power of unifying modeling efforts across research disciplines. Even if BioCRNpyler is not the ideal package for a given system, researchers should strongly consider releasing their models in SBML format, or in a format with sufficient detail to enable conversion to SBML with the txt2biocrnpyler tool or other SBML conversion tools [33]. SBML models allow researchers to quickly reproduce results, expand, or mix and match models to rapidly explore new application spaces. This type of unified modeling approach will be necessary for the field of engineering biology to reach its full potential.

## METHODS

All simulation trajectories presented in this study were generated using deterministic chemical reaction networks; consequently, statistical error bars are not applicable.

The default units for concentration are µmol L^−1^ in BioCRNpyler. However, all examples in this study were conducted using nmol L^−1^ for concentrations of model inputs and model rate constants.

## ASSOCIATED CONTENT

### Code Availability

BioCRNpyler source code and documentation are available as a public repository at https://github.com/BuildACell/bioCRNpyler and https://biocrnpyler.readthedocs.io/en/latest/index.html, re-spectively. Bioscrape source code is available as a public repository at https://github.com/biocircuits/Bioscrape. TMSE-BioCRNpyler Library source code and generated results are available at https://github.com/usnistgov/TMSE_BioCRNpyler_Library. The txt2biocrnpyler source code and generated results are available at https://github.com/usnistgov/txt2biocrnpyler.

## AUTHOR INFORMATION

### Author Contributions

S.W.S. and Z.J. conceived the work, executed the simulations, and drafted and edited the manuscript. Software development was divided between Z.J., who developed the CRN compiler for TMSE circuits, and G.T., who developed the txt2biocrnpyler tool and drafted its associated manuscript sections.

### Funding

Z.J. was supported by the National Research Council Postdoctoral Fellowship. G.T. was supported in part by an appointment to a Research Associateship Program at the National Institute of Standards and Technology, administered by the Johns Hopkins University Whiting School of Engineering.

### Conflicts of Interest

SWS is an inventor on two patent applications pertaining to ctRSD circuits (Application number PCT/US2022/053229) and THE riboregulators (Application number PCT/US25/13610). The authors declare no other conflicts.

### Disclaimer

Certain commercial entities, equipment, or materials may be identified in this document to describe an experimental procedure or concept adequately. Such identification is not intended to imply recommendation or endorsement by the National Institute of Standards and Technology, nor is it intended to imply that the entities, materials, or equipment are necessarily the best available for the purpose. Official contribution of the National Institute of Standards and Technology; not subject to copyright in the United States.

## ACKNOWLEDGMENTS

The authors thank August O. Staubus and Ayush Pandey for insightful feedback on the manuscript. S.W.S also thanks Bright Eyes, whose album The People’s Key resonated as a thematic parallel throughout this work.

## Notes

https://github.com/usnistgov/TMSE_BioCRNpyler_Library

https://github.com/usnistgov/txt2biocrnpyler

### Competing Interest Statement

SWS is an inventor on two patent applications pertaining to ctRSD circuits (Application number PCT/US2022/053229) and THE riboregulators (Application number PCT/US25/13610). The authors declare no other conflicts.

