## Supplementary Material for "Towards interoperable modeling of toehold-mediated strand exchange circuits across DNA nanotechnology and engineering biology"

July 20, 2026

#### Contents

**1 Toehold Mediated Strand Exchange (TMSE) Overview** **S2**
- 1.1 Schematic of key components and TMSE applications . . . . . S2
- 1.2 Schematics of DNA and RNA strands . . . . . S3

**2 The TMSE-BioCRNpyler Library** **S4**
- 2.1 Components . . . . . S4
- 2.2 Mechanisms . . . . . S4

**3 Scaling of class instantiations across modeling abstractions** **S5**

**4 Generating functional BioCRNpyler script using the txt2biocrnpyler** **S5**
- 4.1 Installing and using txt2biocrnpyler . . . . . S5
- 4.2 Required formatting for txt2biocrnpyler . . . . . S6
- 4.3 Expanding literature models via BioCRNpyler . . . . . S6
- 4.4 LLM performance in zero-shot BioCRNpyler script generation . . . . . S7

**5 Validation of TMSE-BioCRNpyler Library** **S7**
- 5.1 Benchmarking against the Jung2022 OR model . . . . . S7
- 5.2 Benchmarking against the ctRSD simulations . . . . . S10

**6 Additional Examples** **S13**
- 6.1 Optional TMSE mechanisms . . . . . S13
- 6.2 Modeling seesaw circuits . . . . . S14
- 6.3 Modeling different thresholding TMSE circuits . . . . . S16
- 6.4 Modeling a compartmentalized DNA-based TMSE circuit . . . . . S17
- 6.5 Modeling a mixed nucleic acid TMSE circuit with different enzyme kinetics . . . . . S18

*Disclaimer*: Certain commercial entities, equipment, or materials may be identified in this document to describe an experimental procedure or concept adequately. Such identification is not intended to imply recommendation or endorsement by the National Institute of Standards and Technology, nor is it intended to imply that the entities, materials, or equipment are necessarily the best available for the purpose. Official contribution of the National Institute of Standards and Technology; not subject to copyright in the United States.

### 1 Toehold Mediated Strand Exchange (TMSE) Overview

#### 1.1 Schematic of key components and TMSE applications

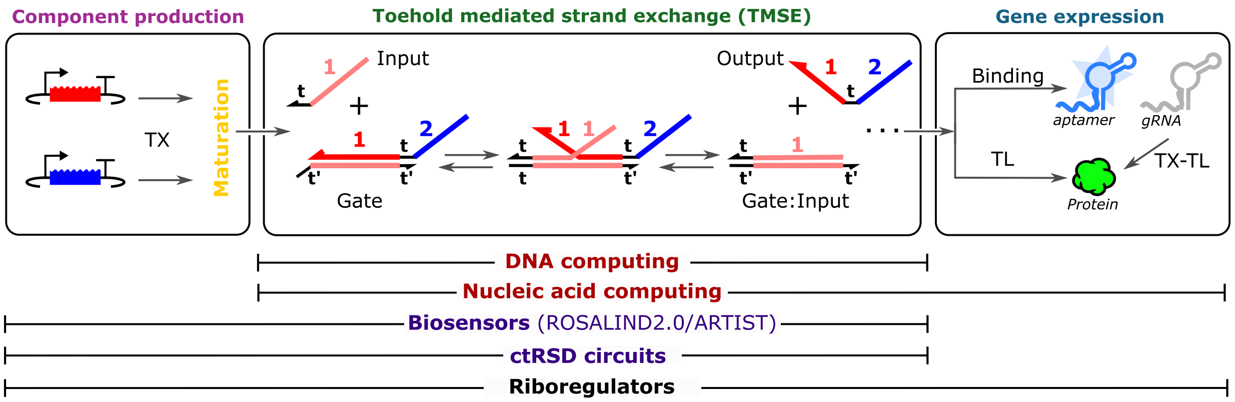


Figure S1: Illustration of the key components and mechanisms added or modified in BioCRNpyler to model TMSE circuits: component production (encompassing RNA maturation), TMSE, and gene expression. Labels underneath indicate prominent applications for TMSE circuits and the modules they span. DNA computing in enzyme-free conditions [1]; Nucleic acid computing in mammalian cells [2, 3]. Biosensors [4, 5, 6]; ctRSD circuits [7, 8]; riboregulators [9, 10, 11].

#### 1.2 Schematics of DNA and RNA strands

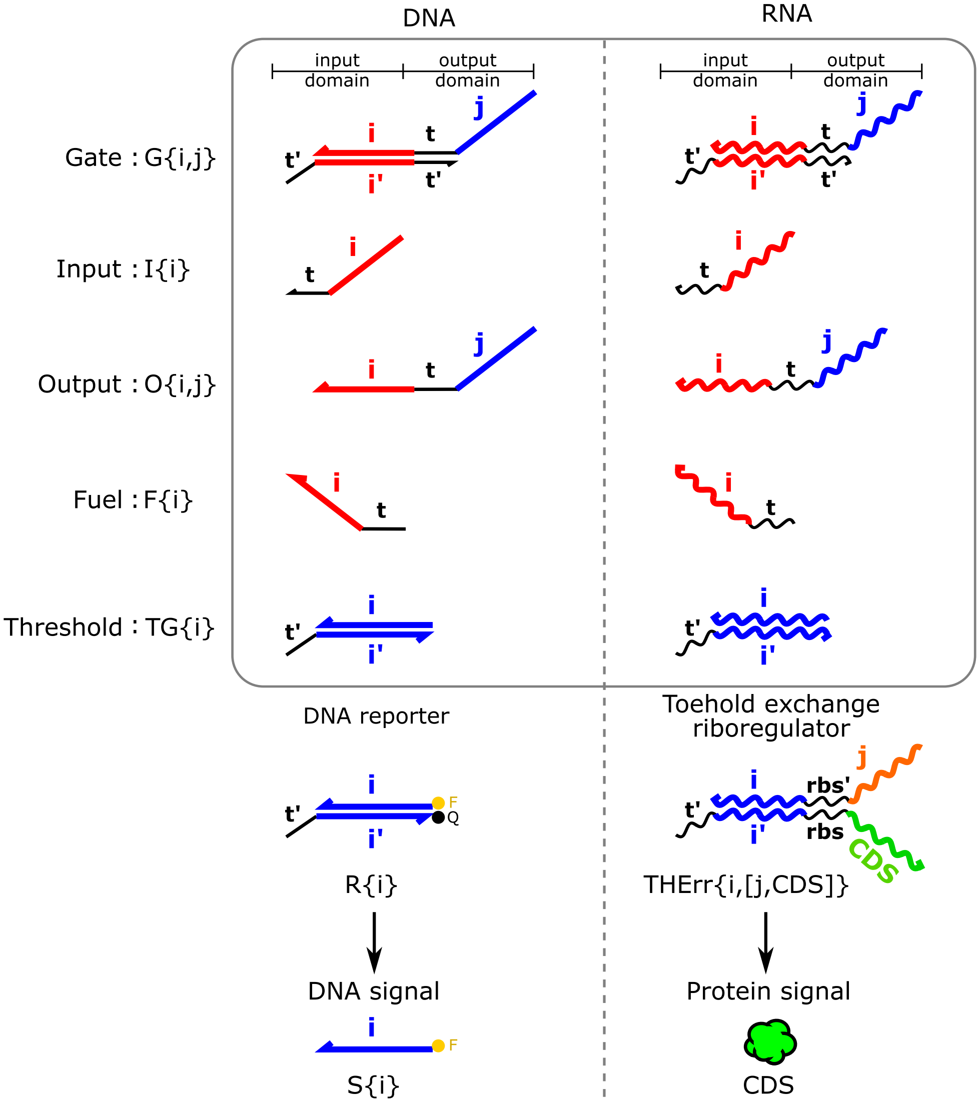


Figure S2: Schematics of DNA (left) and RNA (right) TMSE molecules considered in this study, using the naming convention described in [10] and [12]. *t* specifies a toehold, *i* specifies an input domain, *j* specifies an output domain, *rbs* specifies a ribosome binding site, *CDS* specifies a protein coding sequence. Apostrophes specify sequence complementarity.

### 2 The TMSE-BioCRNpyler Library

#### 2.1 Components

The following subsections provide a list of all new components available for the TMSE-BioCRNpyler tool.

Table S1: TMSE-BioCRNpyler component library.

| Name | Description |
|---|---|
| `LeakctRSD()` | Gate DNA that exhibits leak behavior, allowing for unintended transcription. |
| `StrandExchangeMol()` | Molecules that can undergo a strand exchange. |
| `RNAmaturation()` | RNA molecules that require processing prior to strand-exchange. |
| `TMSE()` | Molecules that undergo two-domain mediated strand exchange. |

The `TMSE()` component module starts by sorting strand exchange molecules into arrays based on the name of the strand exchange molecule — 'Gate', 'TGate', 'Fuel', 'Reporter' and 'Input' — after checking that the given species have valid indices. Strand exchange reactions are appended to the reaction dictionary based on the following logic, strand exchange occurs:

- for $\text{Input}_{i,j}$ and $\text{Gate}_{k,l}$ if j=k
- for $\text{Output}_{i,j}$ and $\text{Gate}_{k,l}$ if j=k
- for $\text{Fuel}_{i,j}$ and $[\text{Input}_{k,l} + \text{Output}_{k,l}]$ if j=k
- for $\text{TGate}_{i,j}$ and $[\text{Input}_{k,l} + \text{Output}_{k,l}]$ then if i=l or j=k
- for $\text{Reporter}_{i,j}$ and $[\text{Input}_{k,l} + \text{Output}_{k,l}]$ then if i=l or j=k

Table S2: New RNA Assembly library component.

| Name | Description |
|---|---|
| `RNAassembly()` | High-level representation of an RNA expression construct. |

#### 2.2 Mechanisms

The following subsections provide a list of all new mechanisms available for the TMSE-BioCRNpyler tool.

Table S3: TMSE-BioCRNpyler mechanism library.

| Name | Description |
|---|---|
| `Maturation01()` | First order maturation mechanism for preprocessed molecules. |
| `GateTxLeak()` | Leakage mechanism for unintended transcription of Gate DNA to Output strands. |
| `OneStepTMSE()` | One step catalytic mechanism for strand exchange. |
| `TwoStepTMSE()` | Two step catalytic mechanism for strand exchange. |
| `ThreeStepTMSE()` | Three step catalytic mechanism for strand exchange. |

### 3 Scaling of class instantiations across modeling abstractions

Table S4: Scaling of class instantiations across different modeling abstractions. The number of `Reaction.from_massaction()` calls required to manually define a circuit grows rapidly with both the number of species and the complexity of the chosen mechanism (`OneStep()`, `TwoStep()`, or `ThreeStep()`), quickly making manual component declarations unwieldy for literature-scale models.

| # of species | OneStep() | TwoStep() | ThreeStep() |
|---|---|---|---|
| 1 | 1 | 2 | 3 |
| 2 | 2 | 4 | 6 |
| 3 | 3 | 6 | 9 |
| 4 | 4 | 8 | 12 |
| ... | ... | ... | ... |
| Ref [13] | 74 | 148 | 222 |
| Ref [14] | 225 | 450 | 675 |
| Ref [15] | > 700 | > 1400 | > 2100 |

### 4 Generating functional BioCRNpyler script using the txt2biocrnpyler

#### 4.1 Installing and using txt2biocrnpyler

Generating a functional BioCRNpyler script or Systems Biology Markup Language (SBML) XML files using the txt2biocrnpyler GUI. The following steps outline how to setup, launch, and use the txt2biocrnpyler browser-based GUI to convert chemical reaction equations into a functional BioCRNpyler script.

1. **Clone the Repository:** Download the tool by cloning the repository at `https://github.com/usnistgov/txt2biocrnpyler`.

2. **Install Dependencies:** Follow the instructions provided in the repository's `README.md` to install all required packages, including Bioscrape (if it is not already installed on your system).

3. **Launch the Interface:** Open the txt2biocrnpyler.ipynb Jupyter Notebook and satisfy the listed prerequisites. Run the notebook cell to launch the txt2biocrnpyler Streamlit GUI, which will open directly in your default web browser.

4. **Upload Source File:** Upload your reaction descriptions as a text file or paste the text directly into the Biochemical text box.
   *Note: In the example below, we imported pages 1 to 4 from the Supporting Information of Jung et al. [4], which details the NOT circuit model utilized in Figure 3, into an AI tool to obtain the reactions in the format detailed in Section 4.2.*

5. **Confirm and Convert:** Review the interface to verify that the contents of your uploaded text file are displayed correctly, then click Convert.

6. **Review Extracted Data:** Once the conversion is complete, navigate through the Species, Reactions, and Parameters tabs to verify the model details are accurate.

7. **Download XML:** Click Download SBML XML to save your final, simulation-ready file.

8. **Download BioCRNpyler:** Click Download BioCRNpyler file to save a prefilled .py file.

**Troubleshooting and Support:** If you encounter any errors during installation or file conversion, please consult the repository's README.md. If the problem persists, please submit an issue on the repository.

#### 4.2 Required formatting for txt2biocrnpyler

##### Chemical reaction format

```
// ------------------------------------------------
// Reactions
// ------------------------------------------------
Reactant_A + Reactant_B --(rate_constant_1)--> Product_C
Reactant_B + Reactant_C <--(rate_constant_2F/rate_constant_2R)--> Product_D + Product_E
Product_C --(rate_constant_3)--> Reactant_A
Product_E --(rate_constant_3)--> Reactant_B

// ------------------------------------------------
// Reaction Rates (Parameters)
// ------------------------------------------------
rate_constant_1 = 10 000.0   // L/umol-s  (Optional description here)
rate_constant_2F = 5000.0    // L/umol-s
rate_constant_2R = 1000.0    // L/umol-s
rate_constant_3 = 0.0003    // 1/s

// Notes on Conservation Laws or calculations:
// Free_Reactant_A = Total_A - Product_C
```

#### 4.3 Expanding literature models via BioCRNpyler

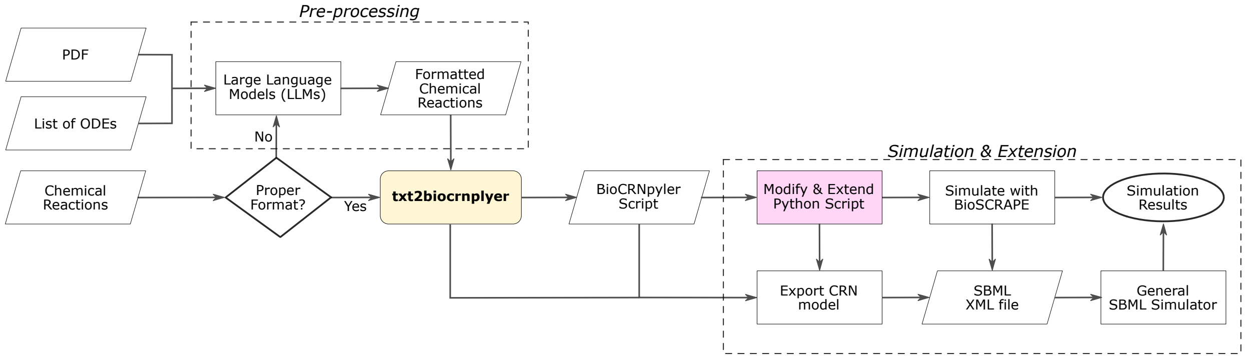


Figure S3: Workflow for automated model generation, extension, and simulation using txt2biocrnpyler. In this flowchart, rectangles represent processing steps or software tools (e.g., txt2biocrnpyler), parallelograms denote input formats, diamonds indicate validation or decision points, and ovals mark the end points of the workflow. In the Pre-processing step, users must convert unstructured inputs — such as literature PDFs, lists of ordinary differential equations (ODEs), or improperly formatted chemical reactions — into standardized, formatted chemical reaction strings using LLMs or manually. Once properly formatted, these chemical reactions are fed directly into the txt2biocrnpyler tool. The txt2biocrnpyler tool parses the chemical reactions and generates a BioCRNpyler Python script. Within the Simulation & Extension step, users can modify and extend this Python script to incorporate additional reactions or contextual complexity. The pipeline offers highly flexible simulation and export pathways: networks can be simulated directly via Bioscrape to generate results. Alternatively, users can bypass Bioscrape and export the initial output, the base Python script, or the extended model directly to a standardized SBML XML file. This XML file can then be shared or simulated by any general SBML simulator to yield final simulation results.

#### 4.4 LLM performance in zero-shot BioCRNpyler script generation

Table S5: Zero-shot performance of LLMs in generating BioCRNpyler scripts for the NOT circuit, as described in the Supplementary Information of [4].

| LLM Platform | Success | Primary Failure Mode | Key Observations |
|---|---|---|---|
| ChatGPT | 4/10 | Used `Reaction()` instead of `Reaction.from_massaction()`. | Lowest reliability; occasionally varied species names. |
| Gemini | 6/10 | Assigned rate constant to k instead of k_forward. | Moderate reliability. |
| Claude | 10/10 | None. (All scripts executable.) | Validated syntax internally; often provided raw SBML XML. Frequently varied species names. |

When the models failed to produce executable code, the errors were largely due to syntax hallucinations specific to the BioCRNpyler library framework. For Gemini, the most common error was incorrectly assigning the forward rate constant to a generic k parameter instead of the required k_forward argument. ChatGPT, on the other hand, frequently failed by attempting to instantiate reactions directly via the base `Reaction()` class rather than correctly utilizing the `Reaction.from_massaction()` method. Furthermore, even among the successfully generated scripts, the LLMs struggled with consistency. Both ChatGPT and Claude exhibited inconsistent species naming conventions across iterations, requiring manual intervention to map the correct initial conditions before simulating the resulting SBML files. However, once these initial conditions were correctly aligned, all successfully compiled SBML files were simulated and verified to accurately reproduce the expected AND gate logic.

### 5 Validation of TMSE-BioCRNpyler Library

#### 5.1 Benchmarking against the Jung2022 OR model

To validate that the TMSE-species modules produced the correct reactions and final model, we compared the simulations of models compiled using TMSE-BioCRNpyler Library against the original examples in the literature. The primary goal of this exercise was to confirm that the TMSE-BioCRNpyler Library, we developed identifies the correct interactions and reactions from the starting molecules. As simulations from the literature may use different numerical solvers than that employed by Bioscrape, we did not expect these results to be identical, but we reasoned that a relative error of $< 1\,\%$ between simulations would correspond to nominally the same underlying models—any differences being attributed to the numerical techniques used to simulate the models. At very early timepoints, when the concentrations of the species in the simulations are very low, the relative error can be substantially $> 1\,\%$, despite the models showing good agreement. Thus, we confined our analysis to $> 5\,\mathrm{min}$ in simulation time.

We investigated how the TMSE-BioCRNpyler Library compared to circuits that combine transcriptional regulation of Input RNAs and downstream TMSE reactions (ROSALIND). With these circuits, the Input strands are RNAs produced by transcription and the Gates are DNAs at fixed concentrations, so the `StrandExchangeMol()` and `TMSE()` components, and `OneStepTMSE()` mechanism were used to compile the models in BioCRNpyler. These models were compared to the results of bottom-up developed models previously released as Jupyter notebooks (ROSALIND) [4]. The ROSALIND model uses reversible binding of transcription factors to model regulation of Input RNAs, so we reinstantiated those reactions using the `Reactions.from_ massaction()` function in BioCRNpyler. The rest of the TMSE reactions were compiled using the TMSE-specific modules we developed. Comparing the simulation results for a two-input OR gate showed relative errors of $< 1 \times 10^{-4}\,\%$ (Figure S4).

BioCRNpyler has predeveloped modules for modeling inducible promoters (`RepressiblePromoter()` component) that use a Hill function approximation of transcriptional regulation. Using these predefined functions greatly reduces the number of function calls needed to build a model connecting transcriptional regulation and TMSE circuits, so we also explored how this model compares to the ROSALIND model.

We found this simpler model still qualitatively captured the relevant dynamics, with relative errors of 10 % to 30 % compared to the more detailed model (Figure S5). This approximation could likely be improved if the Hill equation parameters were explicitly fit.

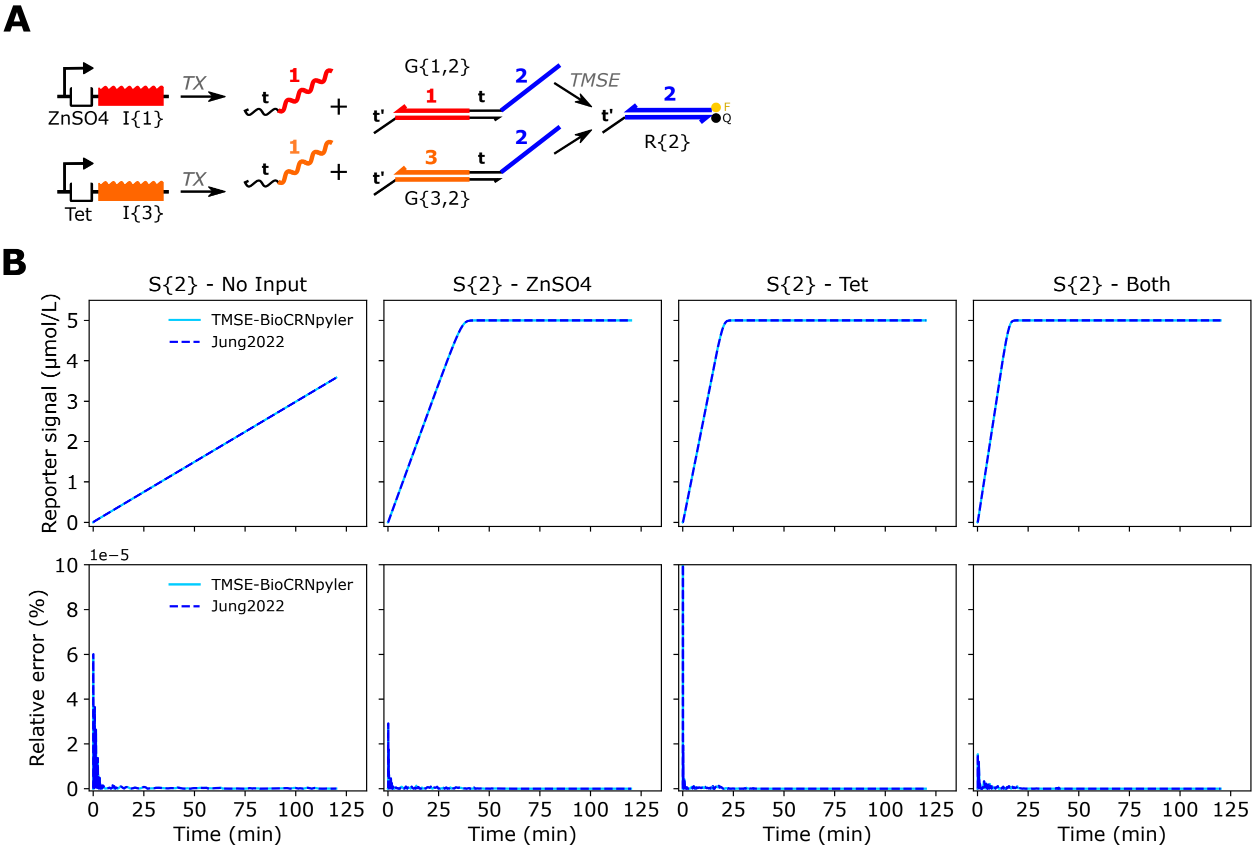


Figure S4: Comparing simulation results of the TMSE-BioCRNpyler Library to previously developed models for ROSALIND [4]. **(A)** Schematic of the two-input OR circuit compared for the two models. **(B)** Simulation results (top row) and relative error (bottom row) for the two models across different inputs. For relative error analysis, the TMSE-BioCRNpyler data considers the Jung2022 results as the "true" value, and the Jung2022 data considers the TMSE-BioCRNpyler results as the "true" value. The Jung2022 model was obtained as a Jupyter notebook from [16].

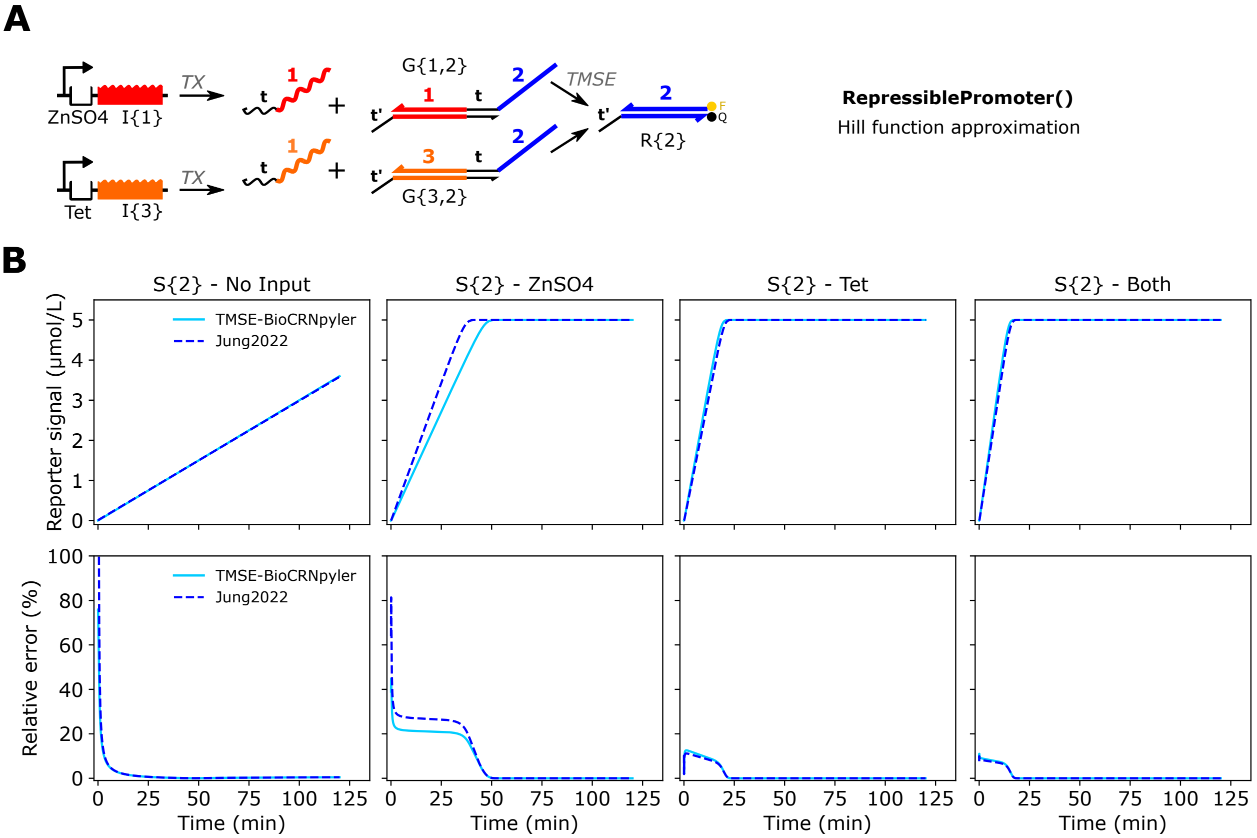


Figure S5: Comparing simulation results of the TMSE-BioCRNpyler Library using the `RepressiblePromoter()` component in BioCRNpyler for transcriptional regulation to previously developed models for ROSALIND [4]. **(A)** Schematic of the two-input OR circuit compared for the two models. **(B)** Simulation results (top row) and relative error (bottom row) for the two models across different inputs. For relative error analysis, the TMSE-BioCRNpyler data considers the Jung2022 results as the "true" value, and the Jung2022 data considers the TMSE-BioCRNpyler results as the "true" value. The Jung2022 model was obtained as a Jupyter notebook from [16]. Using the built-in `RepressiblePromoter()` component in BioCRNpyler allows the system to be modeled using a lumped Hill function instead of the explicit mass-action reactions for reversible transcription factor and polymerase binding used in Figure S4, while still maintaining qualitatively similar results.

#### 5.2 Benchmarking against the ctRSD simulations

Next, we explored how the TMSE-BioCRNpyler Library compared to previously developed models for co-transcriptionally encoded RNA strand displacement (ctRSD) circuits (the ctRSD-simulator) [12]. With these circuits all TMSE components are RNAs produced by transcription so the `StrandExchangeMol()` and `TMSE()` components, and `OneStepTMSE()` and `RNAmaturation()` mechanisms were used to compile the models in BioCRNpyler. These models were then compared to the results from the ctRSD-simulator. Both the TMSE-BioCRNpyler Library and the ctRSD-simulator have options to include ctRSD-specific leak mechanisms, however the implementation of the leak reactions were handled slightly differently between the two packages – with the TMSE-BioCRNpyler Library being more physically accurate. These differences result in the ctRSD-simulator overestimating the concentrations of the Gate and Output species by approximately the leak percentage used. To validate that the TMSE-BioCRNpyler Library models produce the expected interactions and reactions between TMSE components, we set the ctRSD-specific leak percentage to 0 for both models. We simulated three different ctRSD circuits that were previously characterized with the ctRSD-simulator, a two-layer cascade (Figure S6), a single-input and two-output fan-out circuit (Figure S7), and a single-input and single-output Fan-out and Fan-in circuit (Figure S8). Simulation results between the two models showed $< 0.12\,\%$ relative error across the time series for all species, indicating the two models produce nominally the same results.

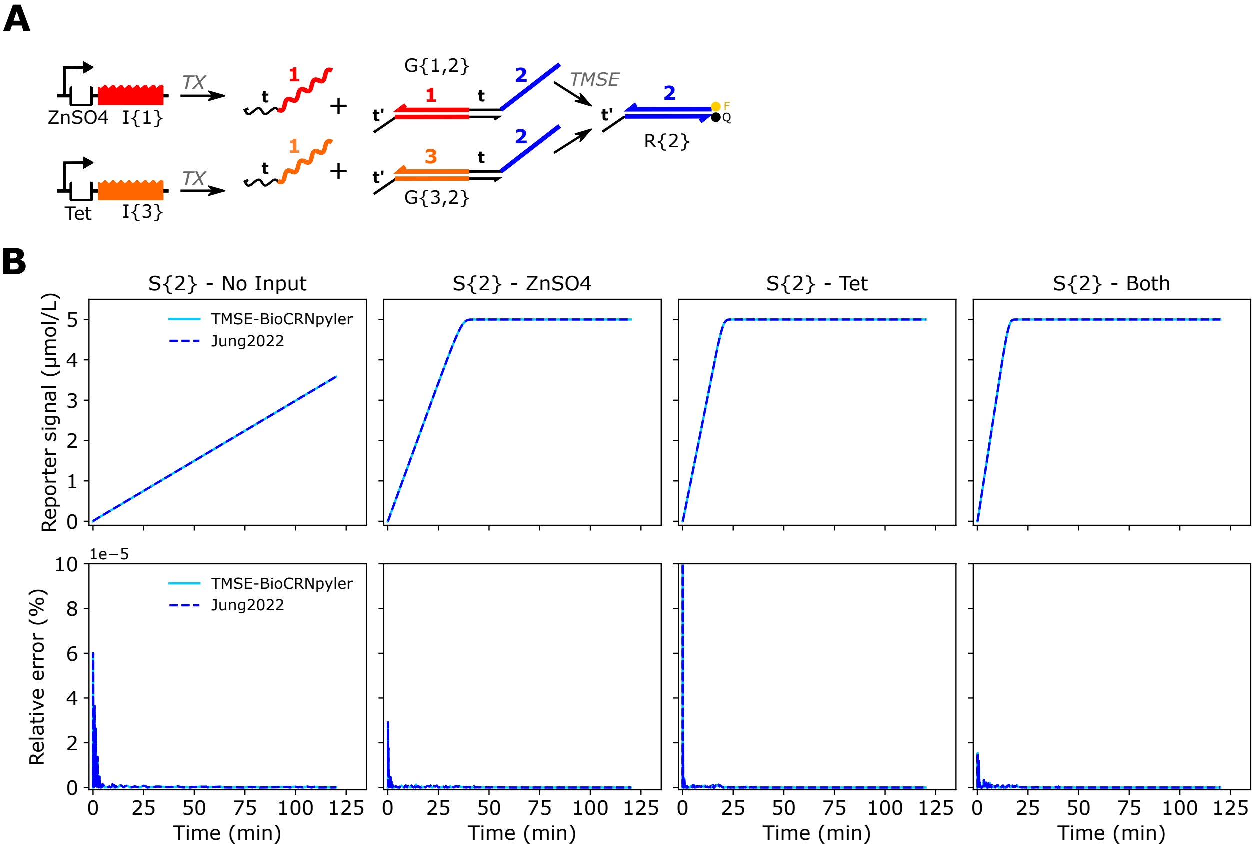


Figure S6: Comparing simulation results of the TMSE-BioCRNpyler Library to the previously developed ctRSD-simulator [17]. **(A)** Schematic of the two-layer cascade compared for the two models. **(B)**Simulation results (top row) and relative error (bottom row) for the two models for different species in the circuit. For relative error analysis, the TMSE-BioCRNpyler data considers the ctRSD-simulator results as the "true" value, and the ctRSD-sim data considers the TMSE-BioCRNpyler results as the "true" value. In the simulations, the concentrations of the I{3}, G{3,1}, and G{1,2} templates were at $25\,\mathrm{nmol\,L^{-1}}$, $25\,\mathrm{nmol\,L^{-1}}$, and $10\,\mathrm{nmol\,L^{-1}}$, respectively. The R{2} reporter concentration was $500\,\mathrm{nmol\,L^{-1}}$. Default ctRSD circuit rate constants were used.

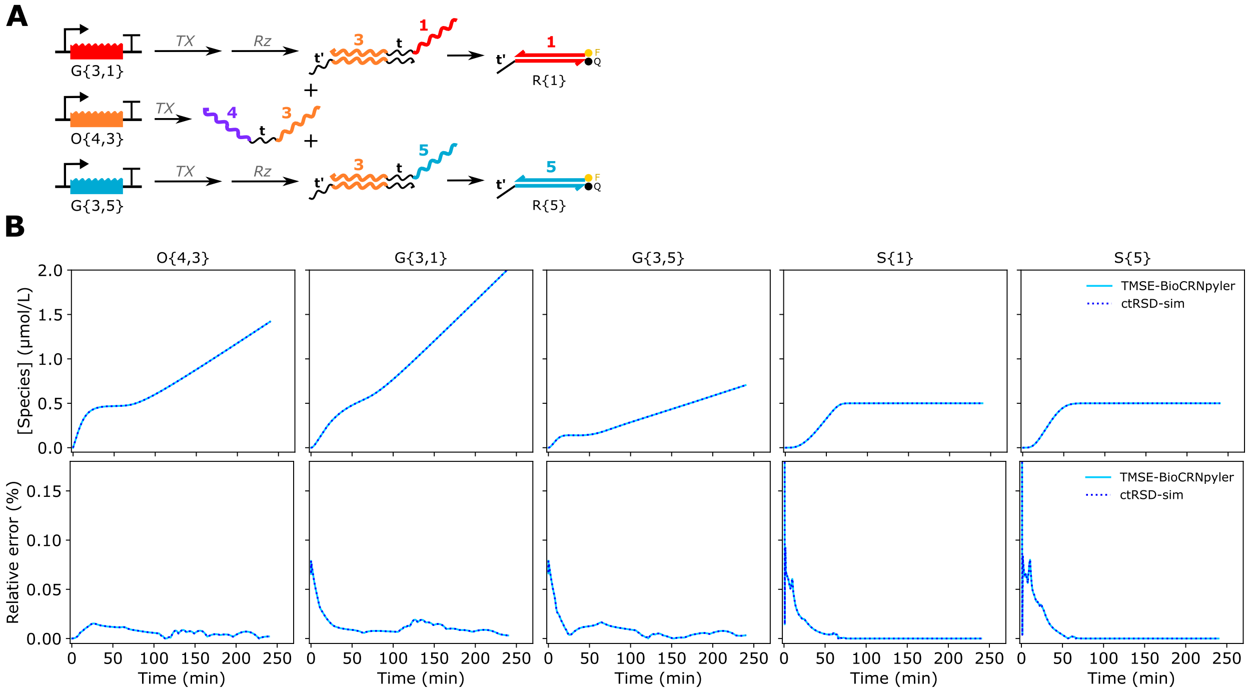


Figure S7: Comparing simulation results of the TMSE-BioCRNpyler Library to the previously developed ctRSD-simulator [17]. **(A)** Schematic of the fan-out circuit compared for the two models. **(B)** Simulation results (top row) and relative error (bottom row) for the two models for different species in the circuit. For relative error analysis, the TMSE-BioCRNpyler data considers the ctRSD-simulator results as the "true" value, and the ctRSD-sim data considers the TMSE-BioCRNpyler results as the "true" value. In the simulations, the concentrations of the O{4,3}, G{3,1}, and G{3,5} templates were at $25\,\mathrm{nmol\,L^{-1}}$, $15\,\mathrm{nmol\,L^{-1}}$, and $15\,\mathrm{nmol\,L^{-1}}$, respectively. Both reporter concentrations were $500\,\mathrm{nmol\,L^{-1}}$. Default ctRSD circuit rate constants were used, except $k_x$ was increased 5-fold for G{3,5} and $k_{rep}$ was increased 5-fold for R{1}.

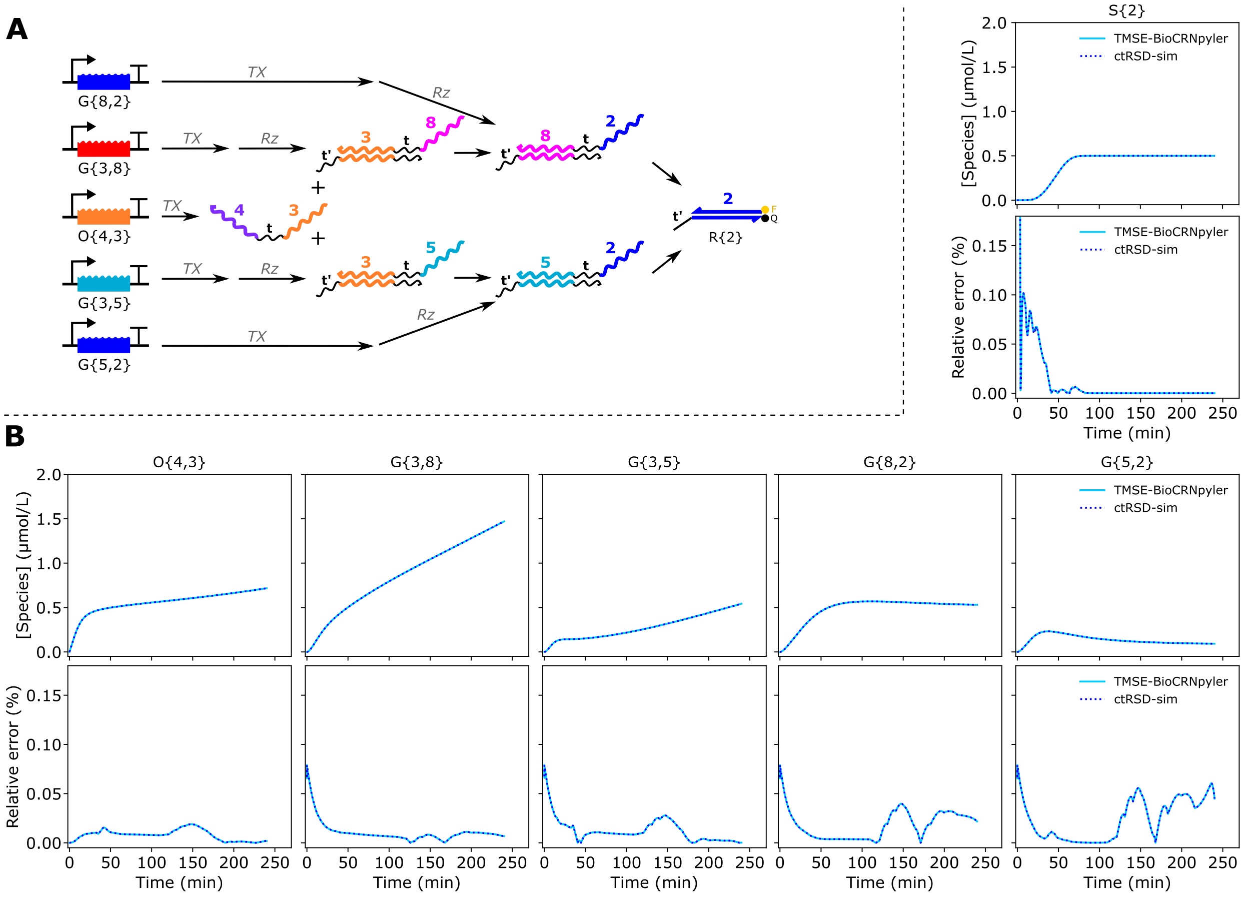


Figure S8: Comparing simulation results of the TMSE-BioCRNpyler Library to the previously developed ctRSD-simulator [17]. **(A)**Schematic of the fan-out / fan-in circuit compared for the two models. **(B)** Simulation results (top row) and relative error (bottom row) for the two models for different species in the circuit. For relative error analysis, the TMSE-BioCRNpyler data considers the ctRSD-simulator results as the "true" value, and the ctRSD-sim data considers the TMSE-BioCRNpyler results as the "true" value. In the simulations, the concentrations of the O{4,3}, G{3,8}, G{3,5}, G{8,2}, and G{5,2} templates were at 25 nmol $L^{-1}$, 15 nmol $L^{-1}$, 15 nmol $L^{-1}$, 10 nmol $L^{-1}$, and 10 nmol $L^{-1}$ respectively. The R{2} reporter concentration was 500 nmol $L^{-1}$. Default ctRSD circuit rate constants were used, except $k_x$ was increased 5-fold for G{3,5} and 3-fold for G{5,2} and $k_{rev}$ was decreased to 10 L $mol^{-1}$ $s^{-1}$ for G{5,2} and G{8,2}.

### 6 Additional Examples

#### 6.1 Optional TMSE mechanisms

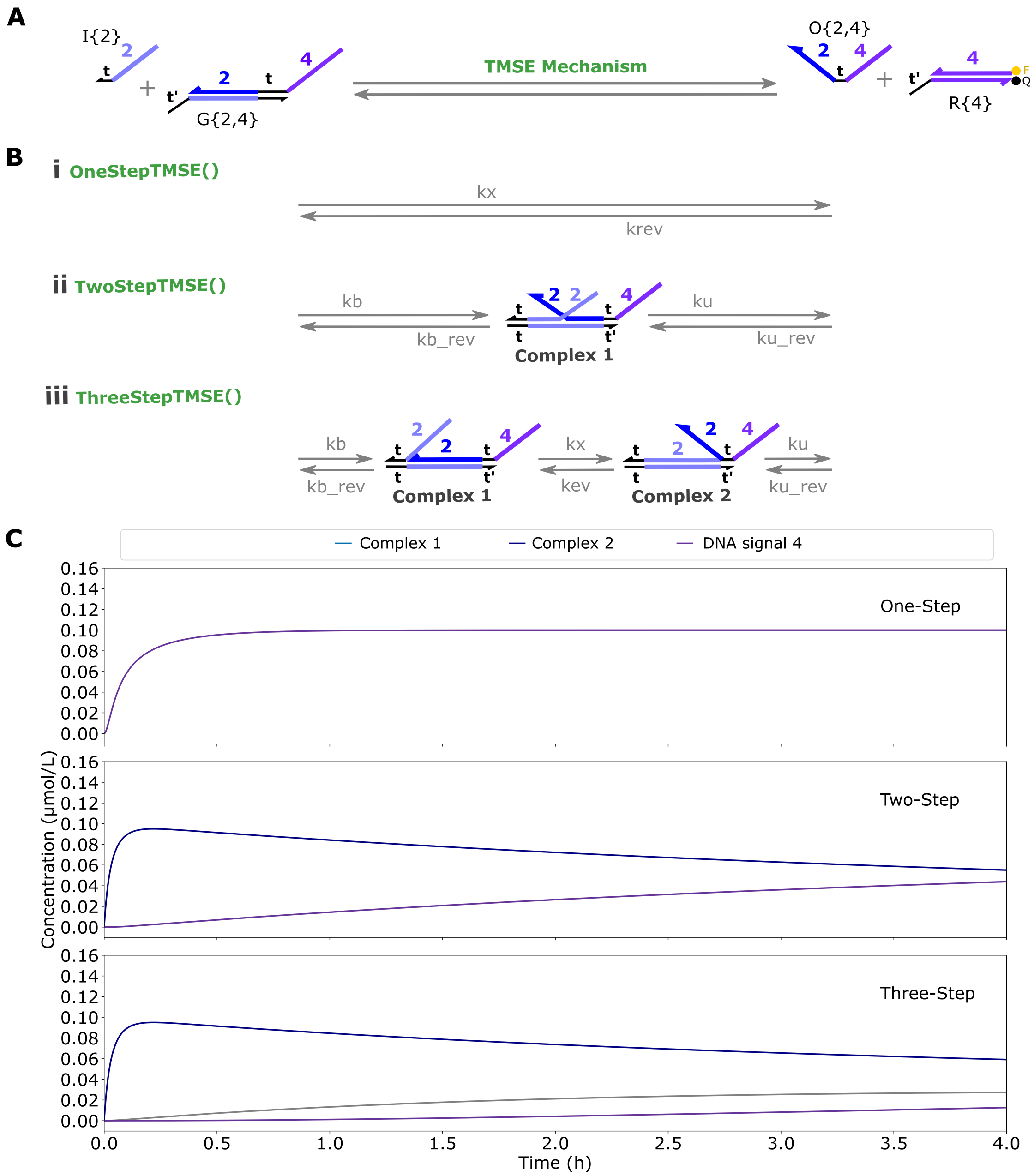


Figure S9: Toehold Mediated Strand Exchange (TMSE) mechanisms. **(A)** Substrates (left) and products (right) involved in the TMSE mechanism. **(B)** Schematic representation of different TMSE mechanisms available: **(i)** `OneStepTMSE()`, **(ii)** `TwoStepTMSE()`, and **(iii)** `ThreeStepTMSE()`. Intermediate species are labeled as Complex 1 and Complex 2. **(C)** Simulation results for each TMSE mechanism, with Complex 1 represented in blue, Complex 2 in gray, and final signal indicated in purple.

#### 6.2 Modeling seesaw circuits

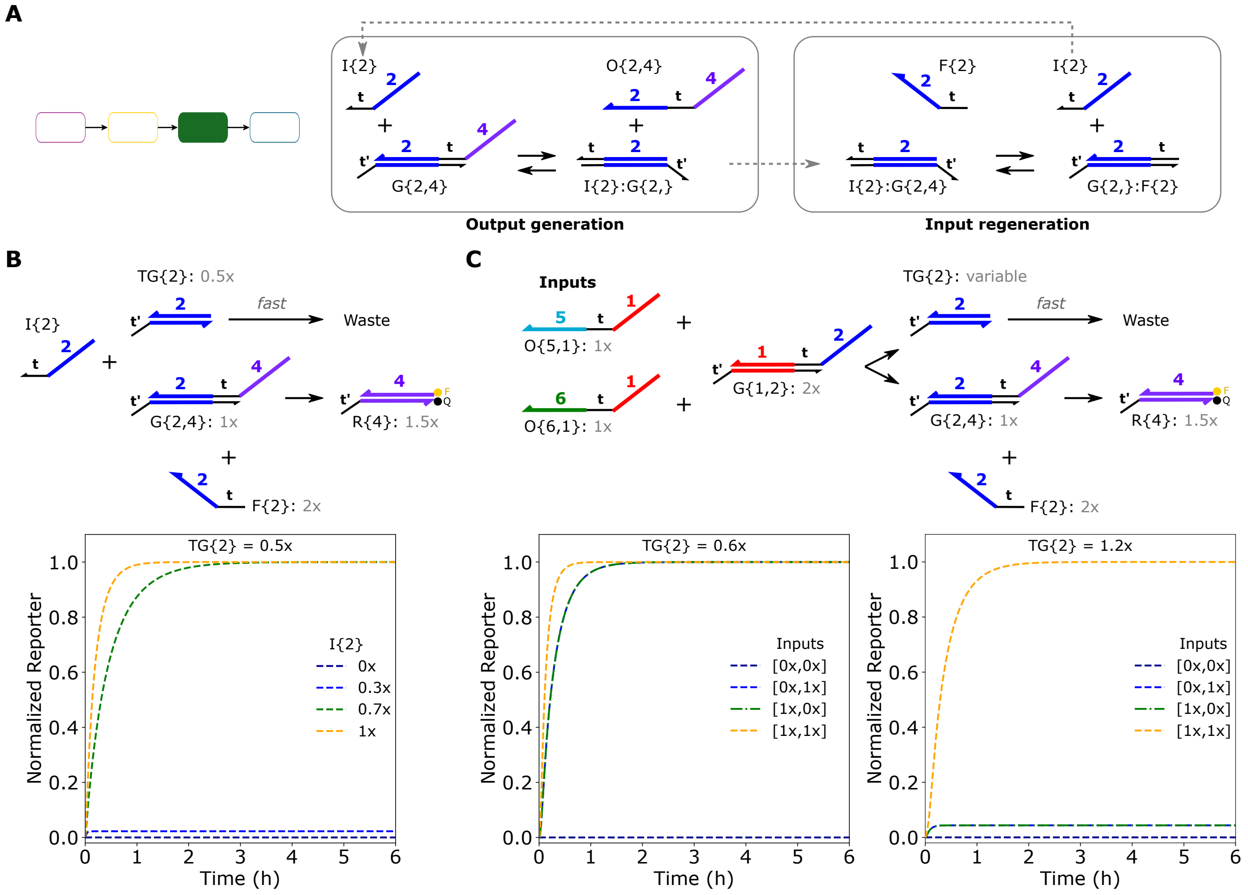


Figure S10: Modeling seesaw circuits from [1] using just the TMSE module. **(A)** Schematic of the seesaw circuit reactions. An Input and Gate react to produce an Output strand (left). This reaction produces an Input:Gate complex with an exposed toehold. This exposed toehold can recruit a Fuel strand to bind and displace the Input strand (right). The Input is then free to initiate another round of Output generation. **(B)** The seesaw reaction is often coupled with a Threshold reaction to enable the logic operations. The Threshold is designed to react with the input much faster than the input reacts with the Gate ($\approx$ 100-fold greater rate constant). So, the signal will only propagate to the reporter if the concentration of the Input is greater than the concentration of the Threshold. The seesaw reaction ensures that 100 % of the Gate is converted to Output by any concentration of Input that exceeds the threshold. **(C)** Combining the seesaw and thresholding reactions enables digital logic operations. OR logic is obtained using a Threshold concentration below either of the two Inputs individually (left). AND logic is obtained using a Threshold concentration that is slightly higher than either Input concentration individually (right). In these simulations the $1\times$ concentration is $100\,\mathrm{nmol\,L^{-1}}$. Colored blocks in (A) indicate which TMSE-specific modules from Figure 2A were used to create the model.

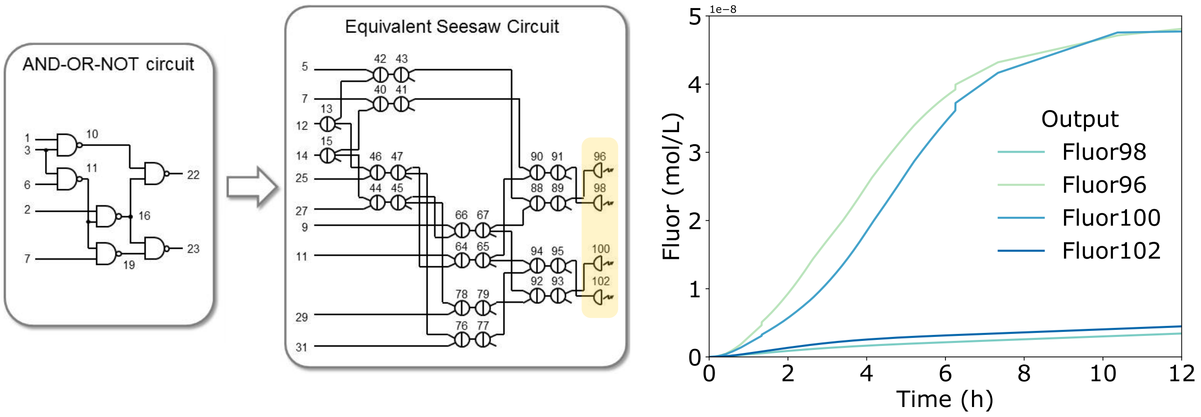


Figure S11: Simulation of an AND-OR-NOT circuit with six NAND gates, consisting of 520 DNA strand exchange molecules, that executes cellular automata transition functions. The circuit schematics (left) and SBML formatted file were obtained from [1].

##### 6.3 Modeling different thresholding TMSE circuits

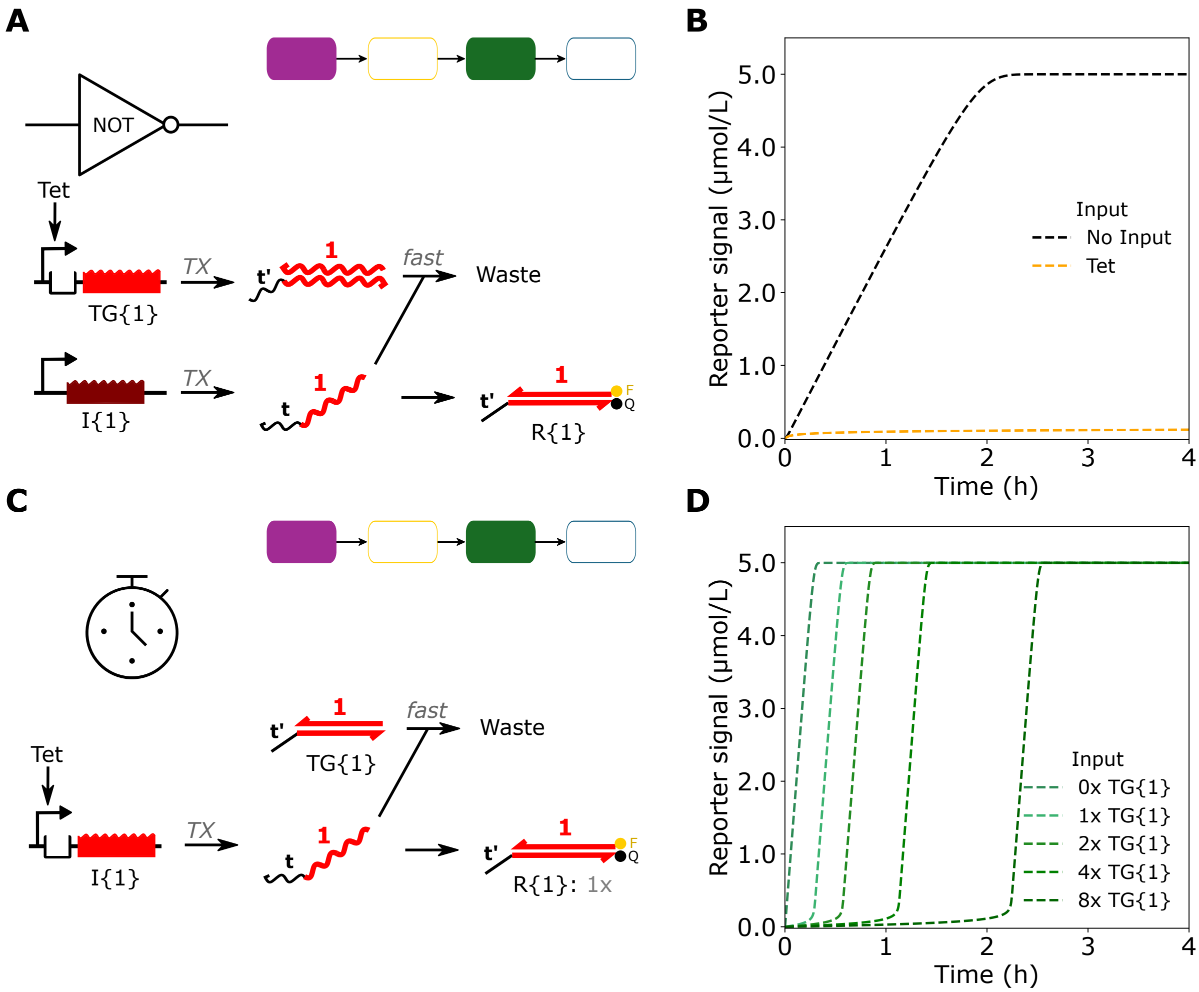


Figure S12: Modeling different thresholding examples from [4]. **(A,B)** Schematic (A) and simulation results (B) of a circuit that executes a NOT operation on a small molecule input (Tet). In [4] the Threshold is designed as a hairpin, which we model as a dsRNA Threshold without an RNA maturation step. **(C,D)** Schematic (C) and simulation results (D) of a circuit that executes a timing operation on a small molecule input (Tet). The DNA Threshold is added at a fixed concentration rather than produced via transcription, such that the more threshold added, the longer it takes the system to produce a measurable signal. Colored blocks above the circuit schematic indicate which TMSE-specific modules from Figure 2A were used to create the model for each simulation.

#### 6.4 Modeling a compartmentalized DNA-based TMSE circuit

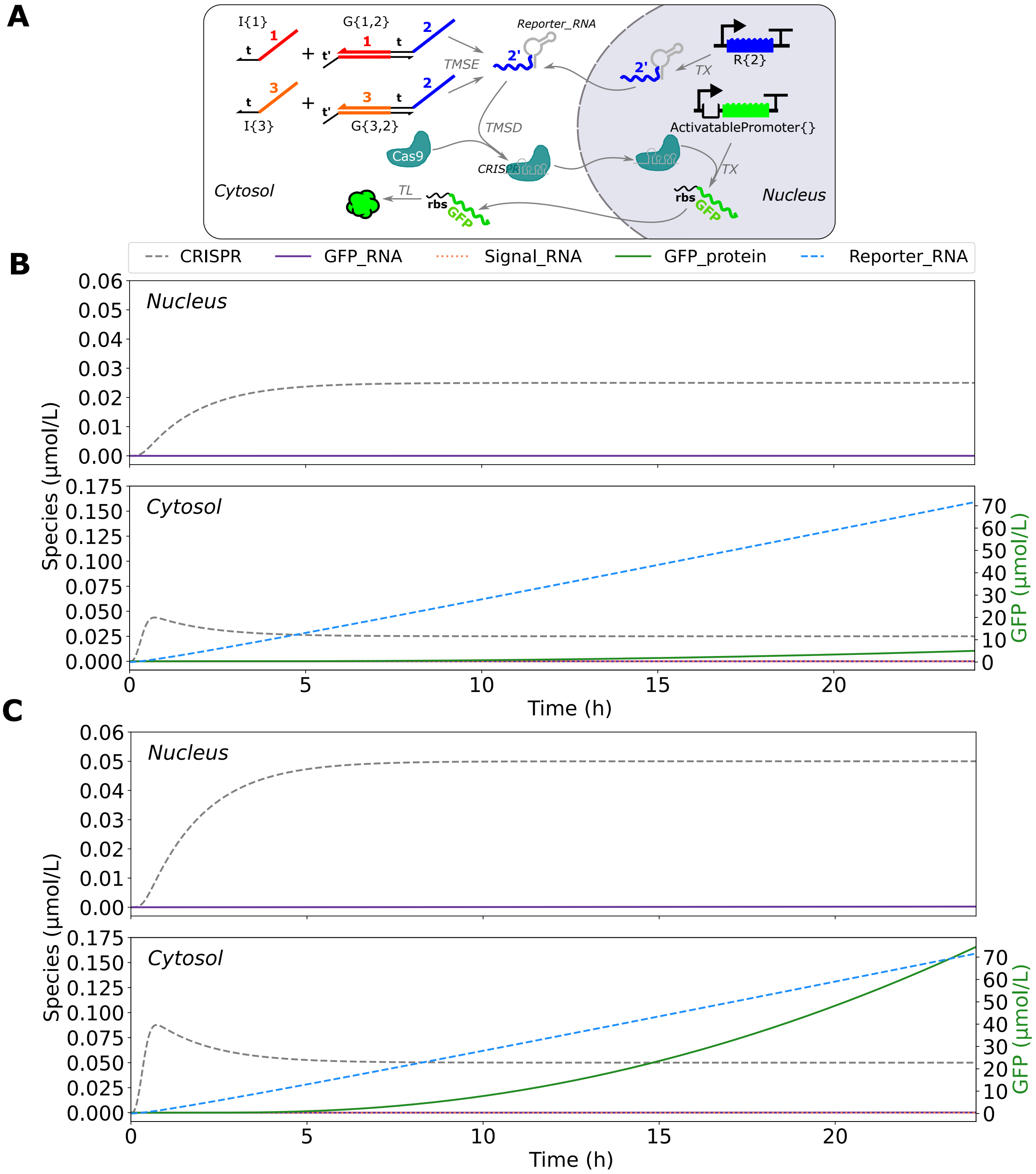


Figure S13: Schematic and simulation of DNA-based TMSE circuits designed to regulate gene expression in mammalian cells with compartmentalization [3]. **(A)** Schematic representation of the nucleus and cytosol compartments, including the respective molecules and their movement between the two cellular regions using the `Mixture()` class in BioCRNpyler. Simulations depict time course measurements of (Top) nucleus and (Bottom) cytosol species when **(B)** either Input 1 or Input 3 is present and **(C)** both Input 1 and Input 3 are present. The green fluorescent protein (GFP) levels are presented on the secondary y-axis.

##### 6.5 Modeling a mixed nucleic acid TMSE circuit with different enzyme kinetics

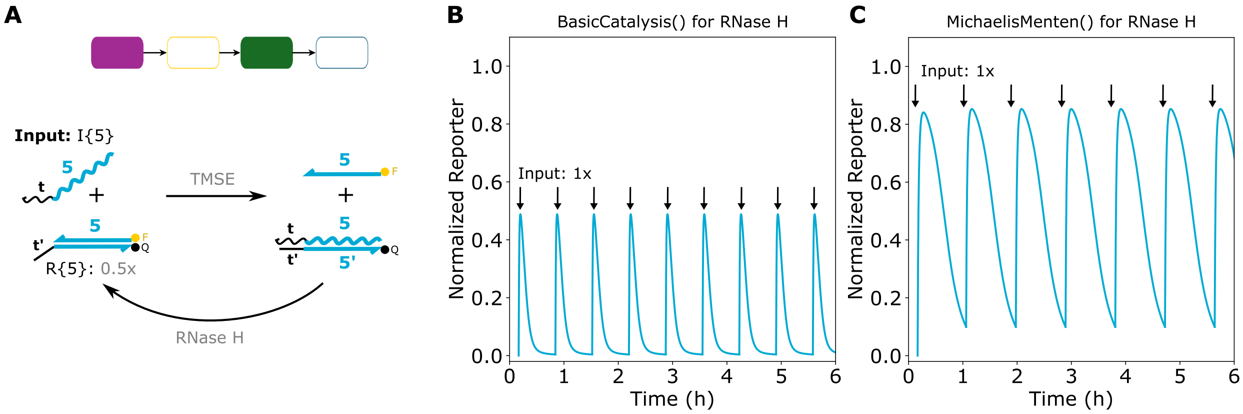


Figure S14: Modeling a mixed nucleic acid system with an added enzyme and discontinuous simulation from [18]. **(A)** Schematic of the system. An RNA Input (I{5}) can react with a Reporter via `TMSE()` to produce a measurable signal. The RNA in the resulting RNA:DNA duplex can be degraded by RNase H to reverse the signal, resulting in a signal pulse. **(B,C)** Simulating the system in panel (A) using BioCRNpyler mechanisms `BasicCatalysis()` or `MichaelisMenten()` for RNase H degradation. In these simulations, 1× is defined as $100\,\mathrm{nmol\,L^{-1}}$. I{5} RNA was spiked in at 1× concentration approximately every hour over the course of the simulation. Colored blocks in (A) indicate which TMSE-specific modules from Figure 2A were used to create the model for the simulation.
